# Affinity requirements for control of synaptic targeting and neuronal cell survival by heterophilic IgSF cell adhesion molecules

**DOI:** 10.1101/2021.02.16.431482

**Authors:** Shuwa Xu, Alina P Sergeeva, Phinikoula S. Katsamba, Seetha Mannepalli, Fabiana Bahna, Jude Bimela, S. L Zipursky, Lawrence Shapiro, Barry Honig, Kai Zinn

## Abstract

Neurons in the developing brain express many different cell adhesion molecules (CAMs) on their surfaces, and CAM interactions are essential for the determination of synaptic connectivity patterns. CAM binding affinities can vary by more than 200-fold, but the significance of affinity differences among CAMs is unknown. Here we provide a systematic characterization of the *in vivo* consequences of altering CAM affinity. Interactions between DIP-α and its binding partners Dpr6 and Dpr10 control synaptic targeting and cell survival for *Drosophila* optic lobe neurons. We generated mutations that change DIP-α::Dpr10 binding affinity and introduced these into the endogenous loci. We show that cell survival and synaptic targeting have different affinity requirements, and that there is a threshold affinity required for targeting. Reducing affinity causes graded loss-of-function phenotypes, while increasing affinity rescues cells that would normally die. Affinity reduction can be compensated for by increasing gene copy number.

## INTRODUCTION

Synapses in the central nervous systems of both vertebrates and invertebrates reside within dense and complex neuropils. During the development of “hard-wired” neural systems such as the *Drosophila* brain, axonal and dendritic processes choose genetically specified synaptic targets within environments where they have access to the surfaces of many non-target neurons. Roger Sperry’s chemoaffinity hypothesis proposed that individual neurons in such systems are labeled by molecules that give them unique identities. The modern version of this hypothesis is that cell adhesion molecule (CAM)-like cell surface proteins (CSPs) expressed on interacting neuronal surfaces bind to each other and trigger downstream events that can cause establishment of synaptic connections between appropriate partners. CAM-like CSPs involved in synaptic targeting in both mammals and *Drosophila* include immunoglobulin superfamily (IgSF) proteins, cadherin superfamily proteins, leucine-rich repeat proteins, and teneurins. Some of these proteins bind homophilically, some have unique heterophilic partners, and still others bind to a variety of partners with different affinities (Honig and Shapiro, 2020; Sanes and Zipursky, 2020).

One interaction network of particular interest was discovered in an *in vitro* “interactome” screen of all ~130 IgSF CSPs in *Drosophila* (Özkan et al., 2013). In this network, the “Dpr-ome”, 21 Dpr (Defective Proboscis Retraction) proteins interact in a complex pattern with 11 DIPs (Dpr Interaction Proteins) (Carrillo et al., 2015; Cosmanescu et al., 2018; Tan et al., 2015). Most DIPs bind to multiple Dprs, and vice versa. Some DIPs and Dprs also bind homophilically. In the pupal brain, neurons expressing a particular DIP are often postsynaptic to neurons expressing a Dpr to which that DIP binds *in vitro* (Carrillo et al., 2015; Cosmanescu et al., 2018; Tan et al., 2015). Loss of these Dprs and DIPs can alter synaptic connectivity and cause neuronal death (reviewed by Sanes & Zipursky 2020) (Ashley et al., 2019; Barish et al., 2018; Bornstein et al., 2019; Carrillo et al., 2015; Cheng et al., 2019; Courgeon and Desplan, 2019; Menon et al., 2019; Venkatasubramanian et al., 2019; Xu et al., 2018; Xu et al., 2019).

Compared to binding between secreted ligands and their receptors, CAM interactions such as those within the Dpr-ome network usually have much lower affinities, with dissociation constants (K_D_s) in the μM range. In the *Drosophila* pupal brain, sequencing studies show that each neuron can express 100 or more different CAM genes (Barish et al., 2018; Konstantinides et al., 2018; Kurmangaliyev et al., 2019, 2020; Li et al., 2020; Özel et al., 2021; Tan et al., 2015). The number in vertebrates is comparable (Sarin et al., 2018). Collectively, these CAM interactions can form stable junctions between cells due to avidity effects. In development, neurons form transient interactions with many other cells during axon/dendrite outgrowth and synaptogenesis. Some transient interactions, such as those with guidepost cells and intermediate targets, are genetically specified and help to determine the correct pattern of synaptic connections. Others probably occur randomly as a consequence of the dense packing of the developing neuropil. Only a small fraction of the interactions experienced by a cell during development are with its final synaptic partners. The ability of a cell to form and break transient interactions may be facilitated by having many different CAMs on its surfaces that bind with low affinity. This allows the cell to manipulate the strength of its adhesive interactions with a particular target by modulating the types and levels of multiple CAMs that have partners on that target. By contrast, if cells interacted with each other *via* a small number of high affinity CAMs, these might have to be completely removed in order to allow a neuron to break a transient interaction with an intermediate target.

Within each CAM family, K_D_s for homophilic and heterophilic binding can vary by as much as 100-fold. For example, the vertebrate clustered protocadherin (Pcdh) genes encode ~60 different isoforms, which interact homophilically with K_D_s varying from ~1 μM to ~100 μM. Type I and type II cadherins interact both homophilically and heterophilically, and their K_D_s can vary over a similar range (reviewed by Shapiro & HOnig 2020). *In vitro* studies have shown that differences in cadherin affinities and expression levels can program the organization of cell clusters and the localization of junctions within collections of tissue culture cells (Brasch et al., 2018; Toda et al., 2018). Similar principles may apply *in vivo* for other CAMs. For example, hair cells and supporting cells in the mouse cochlear epithelia in the auditory system are arranged in checkerboard-like patterns. Each cell type selectively expresses either Nectin-1 or Nectin-3. Heterophilic Nectin-1::Nectin-3 interactions are stronger than homophilic Nectin-1 or Nectin-3 interactions. Removing either protein disrupts the checkerboard arrangement of the two cell types (Togashi et al., 2011). Another example is in the lamina region of the *Drosophila* optic lobe (OL), where different cell types all express a single cadherin, CadN, and the concentric organization of the lamina cartridge is controlled by the relative level of CadN. The L1 and L2 neurons at the center of the cartridge express the highest levels of CadN, and the R cells at the periphery of the cartridge express low levels of CadN. Manipulating the relative levels of CadN expression in different cells can change the relative positions of the neurons within the cartridge and alter its pattern of synapses (Schwabe et al., 2014).

Many studies have analyzed the effects of changing CAM expression levels on synaptic connectivity, but the significance of single protein-protein interaction affinity variation is not well understood. A study by Ozkan, *et. al.* in *C. elegans* examined this issue for the SYG-1 and SYG-2 CAMs. The SYG-1::SYG-2 interaction affects the formation of synapses between the HSN neuron and vulval muscles. HSN presynaptic elements are positioned by complexes between SYG-1 on HSN and SYG-2 on guidepost vulval epithelial cells. *syg-1* null mutations disrupt HSN synapse localization, and this phenotype can be rescued by overexpressing SYG-1 in HSN neurons. SYG-1 mutant proteins with modestly reduced affinity for SYG-2 are impaired in their ability to rescue the *syg-1* null mutant, suggesting that SYG-1::SYG-2 binding affinity is important for function (Ozkan reference).

To determine the role of affinity for a CAM binding pair *in vivo*, it is important to alter both members of the pair and to separate the effects of affinity alterations from those of expression level. Here, we systematically altered the affinity of heterophilic interactions between DIP-α and Dpr10 in a context where both proteins were expressed at normal levels, and examined effects on neuronal wiring during OL development. Binding affinities within the Dpr-ome vary from 1 μM to ~200 μM. DIP-α binds to its two Dpr partners, Dpr6 and Dpr10, with relatively high affinity (K_D_s of 2 μM and 1.36 μM, respectively (Cosmanescu et al., 2018; Sergeeva et al., 2020). DIP-α is expressed in several classes of neurons in the medulla of the pupal OL, including Dm1, Dm4 and Dm12, which arborize in the M1, M3 and M3 medulla layers respectively (Figure 2A). The L3 lamina neuron is presynaptic to both Dm4 and Dm12, and it expresses both Dpr6 and Dpr10 (Davis et al., 2020; Kurmangaliyev et al., 2020; Tan et al., 2015; Xu et al., 2018). The loss of DIP-α::Dpr10 interactions causes several phenotypes during development, including apoptosis-mediated cell loss of Dm1, Dm4 and Dm12, ectopic projections of Dm12 processes to the M8 layer, and alteration of synapse number in Dm12. In addition, overexpression of Dpr10 in the M10 layer of the medulla causes both Dm4 and Dm12 neurons to arborize in M10 (Xu et al., 2018). This provides an ideal system in which to study the significance of affinity for a heterophilic binding pair that controls several developmental processes.

In order to separate the effects of binding affinity alterations from those of expression levels, we introduced a series of designed affinity mutations into the endogenous *DIP-α* and *dpr10* genes. We found that two functions of Dpr10::DIP-α interactions, control of cell survival and of targeting specificity, have different affinity requirements. There is no increase in cell death unless affinity is reduced by 20-fold or more. Remarkably, however, more cells survive than in wild-type when affinity is increased by 2-fold. Synaptic targeting defects are observed when affinity is reduced by ~8-fold (to ~11 μM), and the penetrance of these defects reaches a plateau with a ~20-fold reduction (to ~28 μM). This transition between 11 μM and 28 μM is also observed when targeting is assessed by ectopic expression of Dpr10 in M10. In addition, we observe that adding or subtracting copies of affinity mutant genes affects their phenotypes, indicating that subtle changes in expression level can compensate for alterations in affinity. These results suggest that the affinities between DIP-α and its Dpr partners are finely tuned to allow the correct number of DIP-α expressing neurons to survive and form the appropriate synaptic connections.

## RESULTS

### Generation and selection of DIP-α and Dpr10 mutations that change DIP-α::Dpr10 affinity

DIP-α binds to Dpr6 and Dpr10 with affinities of 2.0 μM and 1.36 μM, respectively. DIP-α also binds to itself with an affinity of 24 μM, and heterophilic and homophilic binding use the same interface residues (Cheng et al., 2019; Cosmanescu et al., 2018; Sergeeva et al., 2020). We developed computational approaches that allowed the design of DIP-α and Dpr10 mutants that changed DIP-α::Dpr10 binding affinity *in vitro* (Sergeeva et al., 2020). To determine how changes in affinity affect neuron-neuron recognition events, we selected a set of DIP-α and Dpr10 mutations for *in vivo* studies based on the following criteria: 1) the mutations should alter binding affinity between DIP-α and Dpr10 in a graded fashion, so as to generate a set of proteins with affinities varying over a wide range; 2) the mutations should not change their specificity for binding to other DIP/Dpr proteins; 3) the mutations should not have strong effects on homophilic binding affinity.

Based on these criteria, we chose DIP-α mutants G74A (K_D_=0.9 μM; when expressed *in vivo*, this mutant is designated as *DIP-α^+2F^*), K81Q (K_D_=31.8 μM, *DIP-α^−20F^*), and K81Q G74S (K_D_=68.0 μM, *DIP-α^−50F^*); and Dpr10 mutants V144K (K_D_=11.3 μM, *dpr10^−8F^*) and Q138D (K_D_=27.6 μM, *dpr10^−20F^*)(Sergeeva et al., 2020). To generate a comparable Dpr10 affinity reduction range as that for DIP-α, we designed an additional *dpr10* mutant, V144K Q142E G99D (K_D_=50.0 μM, *dpr10^−40F^*) (Figures 1B, C). The location of the designed mutations in the DIP-α::Dpr10 interface between N-terminal Ig1 domains is indicated in Figure 1A.

**Figure 1.**
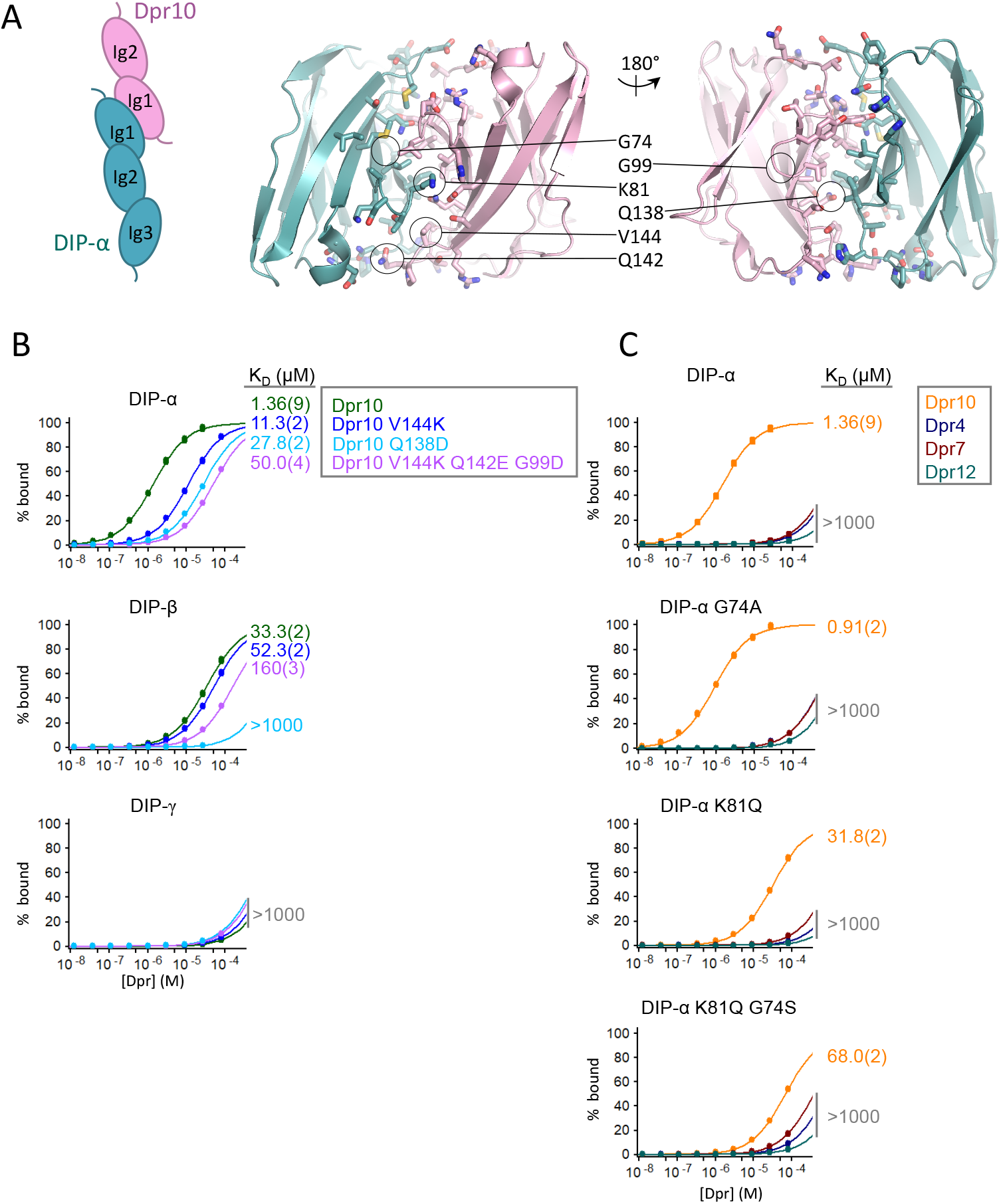
Characterization of DIP-α and Dpr10 affinity mutants. (A) Left: schematic representation of DIP-α/Dpr10 heterodimer formed between N-terminal Ig1 domains of DIP-α (in cyan) and Dpr10 (in pink); Right: the structure of the DIP-α/Dpr10 interface (PDBID: 6NRQ). Interfacial residues (within 6Å of the opposing protomer) are depicted as sticks and mutated positions are encircled and marked. (B, C) Binding isotherms from SPR binding experiments of Dpr10 wild type and its mutants to DIP-α, DIP-β, and DIP-γ (B); and of DIP-α wild type and its mutants to Dpr10, Dpr4, Dpr7 and Dpr12 (C). The binding isotherms and the K_D_s are color-coded according to the legend shown to the right of each panel. K_D_s >1000 μM, describing multiple interactions, are shown in grey. The binding responses for the SPR experiments are shown in Figure S1.

These Dpr10 and DIP-α mutations did not change the specificity of their binding to other DIP/Dpr proteins. Figure 1B shows binding isotherms for interactions of Dpr10 wild-type and mutant proteins with DIP-α, DIP-β and DIP-γ (see Figure S1 for corresponding sensorgrams). DIP-β is closest to DIP-α among all other DIPs in sequence, and is also a Dpr10 binding protein, but with a much weaker affinity (K_D_=33μM). DIP-γ is not a Dpr10 binding protein (K_D_ of >1000 μM). Like wild-type Dpr10, all three mutant Dpr10 proteins interact more strongly with DIP-α than with DIP-β (Figure 1B). None of the mutants displays detectable binding to DIP-γ (Figure 1B). Figure 1C shows binding isotherms for DIP-α wild-type and mutant proteins to Dpr10, Dpr4, Dpr7 and Dpr12 (see Figure S1 for corresponding sensorgrams). Like wild-type DIP-α, none of the DIP-α mutants exhibits measurable binding to Dpr4, Dpr7, or Dpr12, which are members of non-cognate Dpr subgroups (Figure 1C).

Dpr10 is a monomer, while DIP-α is a dimer with a K_D_ of 23.9 μM (Cosmanescu et al., 2018). The DIP-α/DIP-α and Dpr10/DIP-α interfaces are very similar (RMSD of 0.6Å), and hence changes in the heterophilic interface would be expected to change the homophilic DIP-α/DIP-α interface. To ensure that the DIP-α mutants retained the ability to homodimerize, we measured homophilic binding affinities of all DIP-α mutants using analytical ultracentrifugation (AUC). The homophilic binding K_D_ for the three DIP-α mutant proteins are: DIP-α G74A (*DIP-α^+2F^, in vivo*), K_D_=50μM; DIP-α K81Q (*DIP-α^−20F^*) K_D_=19.6μM; DIP-α K81Q G74S (*DIP-α^−50F^*) K_D_=46μM. We confirmed that all mutants remain dimeric, and the changes of homophilic binding between DIP-α in all mutants are no more than 2-fold (Table S1). In summary, these results indicate that we have successfully created Dpr10 and DIP-α mutants with a wide affinity range that do not affect cognate binding preferences of DIPs and Dprs relative to non-cognate partners.

### DIP-α and Dpr10 affinity mutants are expressed normally in vivo

We introduced the chosen mutations into the endogenous *DIP-α* and *dpr10* genomic loci by a precise CRISPR mediated knock-in strategy (Zhang et al., 2014), and sequence verified the generated transgenic animals. We tested expression of the mutant proteins *in vivo* using antibodies specific for DIP-α and Dpr10 (Figures 2B, 2C) (Xu et al., 2018).

**Figure 2:**
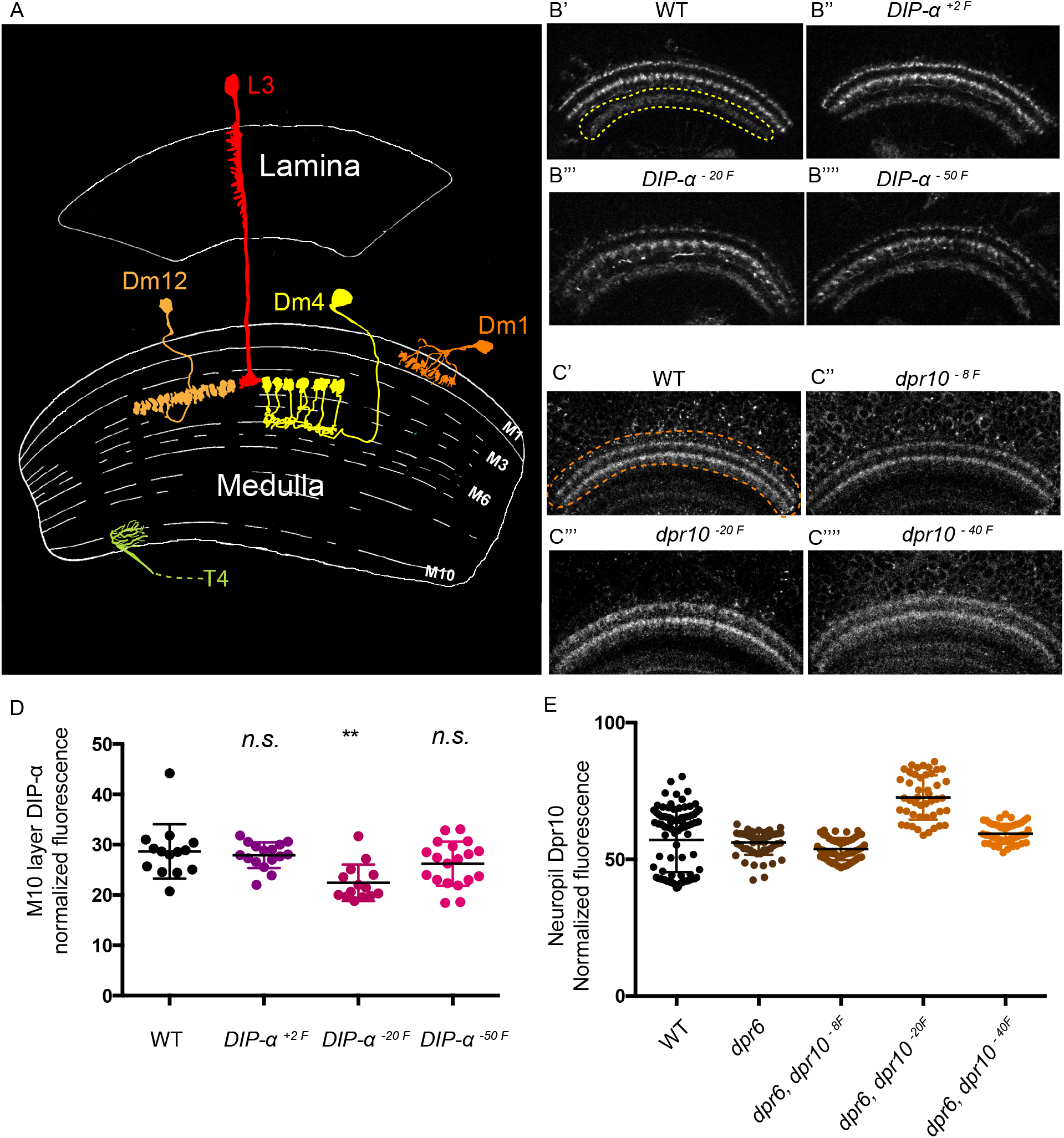
DIP-α and Dpr10 mutant proteins are expressed in wild-type patterns and at comparable levels to wild-type. A) Schematic of the *Drosophila* OL, focusing on the lamina and medulla neuropil areas. Cell types studied in this research are shown. B) Anti-DIP-α antibody staining of medulla region in wild type and *DIP-α* affinity mutants at 48h APF. C) Anti-Dpr10 antibody staining of medulla region in wild type and *dpr10* affinity mutants at 48h APF, D) Quantification of anti-DIP-α fluorescence signal intensity of the 3^rd^ medulla expressing layer at 48h APF (yellow circle lines in B’) in wild type and *DIP-α* affinity mutants. Fluorescence intensity was normalized against background signal. E) Quantification of anti-Dpr10 fluorescence signal intensity of the two Dpr10 expressing neuropil layers (orange dotted circle in C’) in wild type and *dpr10* affinity mutants. Fluorescence intensity was normalized against background signal.

Wild type DIP-α is expressed in three neuropil layers in the medulla region of the OL during mid-pupal development (48h after puparium formation/APF) (Xu et al., 2018). All DIP-α affinity mutants were expressed in the same pattern as the wild type (Figures 2B”-B””). Changing cell surface protein sequences sometimes causes proteins to fail to transport to neuronal terminals, but all of our mutants localized to the medulla neuropil, where axonal and dendritic endings are located (Figures 2B”-B”’).

The DIP-α-expressing Dm1 neuron, and Dm4 and Dm12 neurons, project to the first (M1) and second (M3) DIP-α expressing layers at 48h APF, respectively (Figure 2B’). A large fraction of these neurons undergo cell death during pupal development in *DIP-α* null mutants (see below) (Xu et al., 2018). If our introduced mutations were loss-of-function (LOF), they would be expected to cause a reduction in staining intensity in the first two layers, even if they do not alter the levels of expression in individual cells. Therefore, since another set of DIP-α expressing neurons that project to the third layer (M10; yellow dotted line) do not exhibit detectable cell death in null mutants, we quantitated the expression levels of mutant DIP-α in this layer. All three alleles showed similar expression levels to wild-type DIP-α in this layer (Figure 2D). When anti-DIP-α staining was quantitated across the whole neuropil, DIP-α^+2F^ was present at similar levels as wild-type. Staining intensity for DIP-α^−20F^ and DIP-α^−50F^ was lower than for wildtype, consistent with the idea that some DIP-α-expressing neurons undergo cell death in loss-of-function mutants (Figure S2, and see below).

Dpr10 is expressed in two major medulla layers in the 48h APF optic lobe (Figures 2C’-C””). All three Dpr10 affinity mutant proteins were expressed at the same patterns as the wild type, showing no defects in neuronal terminal localization. Since no Dpr10-expressing OL neurons are known to exhibit cell death, we quantified Dpr10 expression levels in the whole neuropil. Two Dpr10 alleles were expressed at the same level as the wild-type (Dpr10^−8F^ and Dpr10^−40F^), while one was expressed at slightly higher level than wild-type (Dpr10^−20F^).

### Reducing DIP-α::Dpr10 affinity causes graded mistargeting of Dm12 neurons

Null *DIP-α* mutations, or *dpr6, dpr10* double mutations, cause disruption of several cellular processes during OL development, including mistargeting of Dm12 neurons to M8 layer, and death of Dm4 and Dm12 neurons (Xu et al., 2018). To determine how changing binding affinity between DIP-α and Dpr10 affects these processes, we first analyzed neuronal targeting in Dm12 neurons. Dm12 neurons arborize in the M3 layer (Figure 2A). We have previously reported that *DIP-α* null Dm12 clones in a wild-type background target to a more proximal medulla layer, M8 (Xu et al., 2018). We have now developed methodologies to quantitate mistargeting in whole-animal mutants, allowing us to examine much larger numbers of neurons. In *DIP-α* null whole-animal mutants, about one-third (43) of Dm12 neurons mistargeted to M8 (Figures 3E, F). In *DIP-α^−20F^*, which has a ~20-fold reduction in DIP-α::Dpr10 binding affinity, ~5 (4%) Dm12 neurons per OL mistargeted to M8 (Figures 3C, 3F, 3F inset). This number increased to ~20 (17%) in *DIP-α^50F^*, which reduces affinity by ~50 fold (Figure 3D, 3F). The mutation that increased DIP-α/Dpr10 binding affinity by 2 fold (DIP-α^+2F^) produced no change in Dm12 targeting (Figure 3B, 3F).

**Figure 3:**
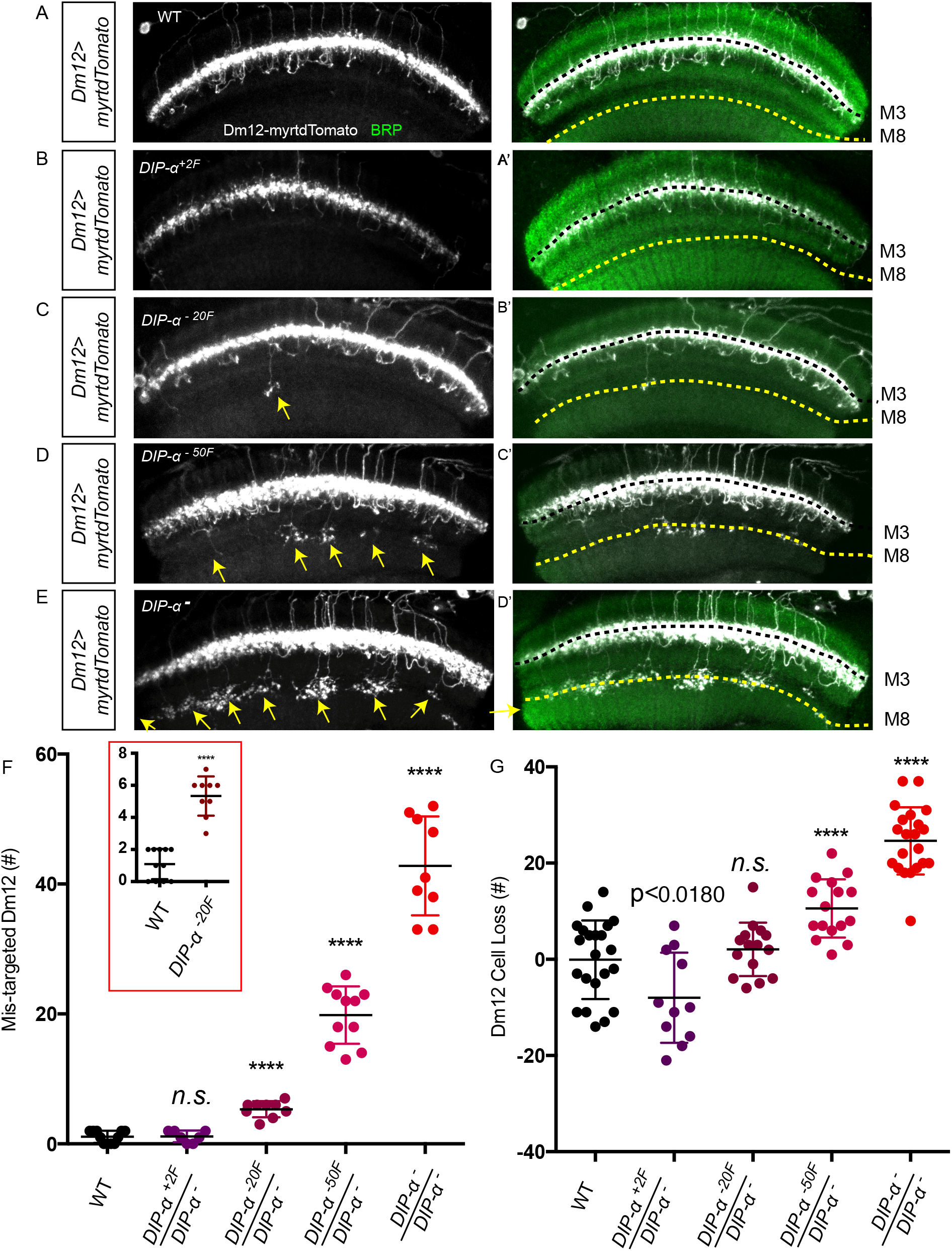
*DIP-α* affinity mutants show different thresholds for synaptic targeting and for cell survival in Dm12 neurons. A-E) Dm12 neurons in wild type, *DIP-α^+2F^/DIP-α^−^, DIP-α^−20F^/DIP-α^−^, DIP-α^−50F^/DIP-α^−^*, and *DIP-α^−^/DIP-α^−^* (null) adult medulla. White, Dm12 neuron labeled with Dm12-LexA>LexAop-myrtdTomato; Green, BRP. Each panel is a maximum intensity projectin (MIP) of a ~10 micron Z stack, 0.18 micron slice interval. Black dotted line: M3 layer; yellow dotted line: M8 layer. F) The number of Dm12 neurons that mistarget to the M8 layer increases as DIP-α::Dpr10 affinity is reduced in *DIP-α* mutants, with significant effects observed for mutants with more than 20-fold reduction (inset). G) The number of missing Dm12 cells also increases as the affinity of DIP-α::Dpr10 is reduced in *DIP-α* mutants, but no significant increase is observed until a 50-fold reduction of affinity is reached.

### Dm12 cell death and mistargeting are differentially affected by DIP-α affinity mutations

In *DIP-α* null mutants, about 25 Dm12 neurons (~22%) die during development, reducing the total Dm12 complement to ~90 (Figure 3F) (Xu et al., 2018). We observed that reducing DIP-α::Dpr10 binding affinity by ~20 fold did not cause any cell loss (*DIP-α^−20F^*). A small number of Dm12 neurons were lost (~8, 5%), when affinity was reduced by 50-fold in *DIP-α^−50F^* mutants (Figure 3G). These results indicate that different affinity thresholds control Dm12 targeting and cell survival. Both the *DIP-α^−20F^* and *DIP-α^−50F^* mutations cause substantial mistargeting but have little or no effect on cell survival (Figures 3E, 3F). The mutation that increased DIP-α/Dpr10 binding affinity by 2 fold (DIP-α^+2F^) produced a slight increase of Dm12 cell number (~7) with p value <0.018 (Figure 3G).

### Differential effects on targeting and cell survival are also observed for *dpr10* affinity mutants

DIP-α binds to Dpr6 and Dpr10 with similar affinities. Both of these Dprs are expressed in the L3 lamina neuron, which forms synapses with the DIP-α-expressing neurons Dm12 and Dm4. Loss of both Dpr6 and Dpr10 (in a double-null mutant) causes cell death for Dm12, Dm4 and Dm1 neurons. *dpr10* or *dpr6* single mutations have milder to no phenotypes, indicating that these Dprs have partially redundant functions (Xu et al., 2018). Thus, to facilitate the analysis of the relationships between Dpr10 affinity and function, we knocked the *dpr10* affinity mutations into the endogenous *dpr10* locus in a *dpr6* null mutant background.

We analyzed Dm12 neurons in the three *dpr10* affinity mutants described above (*dpr10^−8F^, dpr10^−20F^*, and *dpr10^−40F^*). Animals expressing only wild-type Dpr10, but not Dpr6 (*dpr6* single mutant) displayed a mild mistargeting phenotype in Dm12 neurons, with on average ~8 mistargeted Dm12 neurons per OL (Figures 4B, 4G). When Dpr10 affinity to DIP-α was reduced by 8-fold (in *dpr6 dpr10^−8F^*), ~14 mistargeted cells were observed per OL (Figure 4C). A further reduction in affinity to 20-fold less than the wild-type (*dpr6 dpr10^−20F^*) caused a doubling of the number of mistargeted neurons, to ~30 cells per OL (Figure 4D). No further increase in mistargeting was seen in the *dpr6 dpr10^−40F^* mutant, in which affinity is reduced by ~40 fold (Figure 4E). In the double-null mutant, a slightly stronger phenotype was observed, with ~38 Dm12 neurons mistargeting to M8 (Figures 4F, G). Thus, as we showed for *DIP-α* affinity mutants, with the reduction of *DIP-α::Dpr10* affinity, more Dm12 neurons fail to target to the correct layer. However, this change is not linear: there was a major increase in mistargeting when affinity was reduced from 8-fold weaker than wild-type to 20-fold weaker, while reduction from wildtype to 8-fold weaker and from 20-fold weaker to null produced smaller effects (Figure 4G and inset).

**Figure 4:**
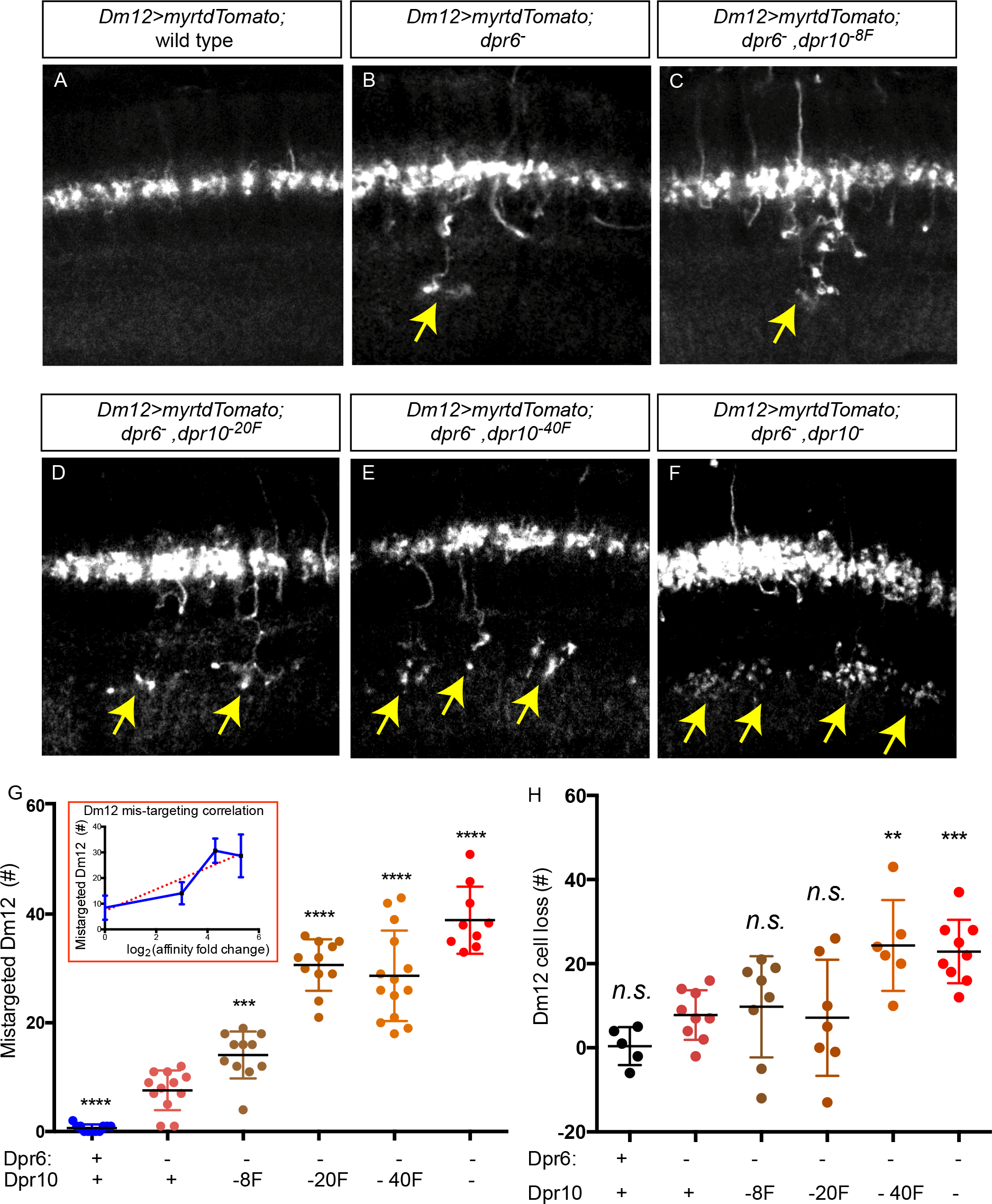
Analysis of *dpr10* affinity mutants shows that there are different affinity thresholds for Dm12 mistargeting and survival. A-F) Images of Dm12 neurons in Dpr10 mutant adult medulla, showing different degrees of mistargeting in the affinity and null alleles (A. WT, B. *dpr6^−^*, C. *dpr6^−^ dpr10^−8F^*, D. *dpr6^−^ dpr10^−20F^*, E. *dpr6^−^ dpr10^−40F^*, F. *dpr6^−^ dpr10^−^*). Each panel is a MIP of a ~10 micron Z stack, 0.18 micron slice interval. Images show representative windows of the entire medulla. G) Increase in mistargeting of Dm12 neurons as affinity between DIP-α and Dpr10 is reduced in *dpr10* mutants. Inset: non-linear relationship between affinity fold decrease and number of mistargeted Dm12 cells. H) Corresponding increase of Dm12 neuron cell loss as affinity between DIP-α and Dpr10 is reduced in *dpr10* mutants.

We also assessed Dm12 cell survival in the affinity mutants. As we previously reported, there was no significant difference in Dm12 numbers between wild-type and *dpr6* single mutants (Xu et al., 2018). No cell loss was seen in the two genotypes bearing mutations that reduce DIP-α::Dpr10 affinity by 8-fold and 20-fold (*dpr6 dpr10^−8F^, dpr6 dpr10^−20F^*).

However, ~24 Dm12 neurons (21%) were lost in the mutant that reduced affinity by 40-fold *(dpr6 dpr10^−40F^*), comparable to the null allele (20%) (Figure 4H).

### An affinity threshold for induction of Dm neuron mistargeting by ectopic Dpr10

The analysis of endogenous *dpr10* knock-in mutants showed that proper affinity between DIP-α and Dpr10 is required for the Dm12 neurons to target to the correct synaptic layer. In addition, a reduction in binding affinity from ~8-fold weaker to ~20-fold weaker than in wild-type produces a major increase in mistargeting. To further evaluate this relationship between affinity and targeting, we used another assay: mistargeting induced by ectopic expression of Dpr10. We previously showed that when Dpr10 is misexpressed at high levels in the M10 layer of the medulla (in T4 cells), the DIP-α expressing neurons Dm4 and Dm12 send their processes to M10, bypassing their normal synaptic layer M3. (T4 is not a synaptic partner of Dm4 or Dm12, but T4 processes come into contact with neurons that project to other medulla layers during early pupal development (Xu et al., 2018). Both Dm4 and Dm12 neurons display mis-expression-induced mistargeting. Because of Dm4’s smaller cell number (40 Dm4s vs 115 Dm12s) and thicker mistargeting axon branch, it is easier to quantitatively assess Dm4 mistargeting events. Thus, we used Dm4 neurons for this analysis. We expressed each of the Dpr10 affinity mutants in M10 using T4-Gal4. Dpr10 antibody staining revealed that M10 layer expression of UAS-Dpr10^−8F^ and UAS-Dpr10^−20F^ is comparable to that of UAS-Dpr10^WT^, while UAS-Dpr10^−40F^ is expressed at 1.6-fold higher levels than wild-type (Figure 5G).

**Figure 5:**
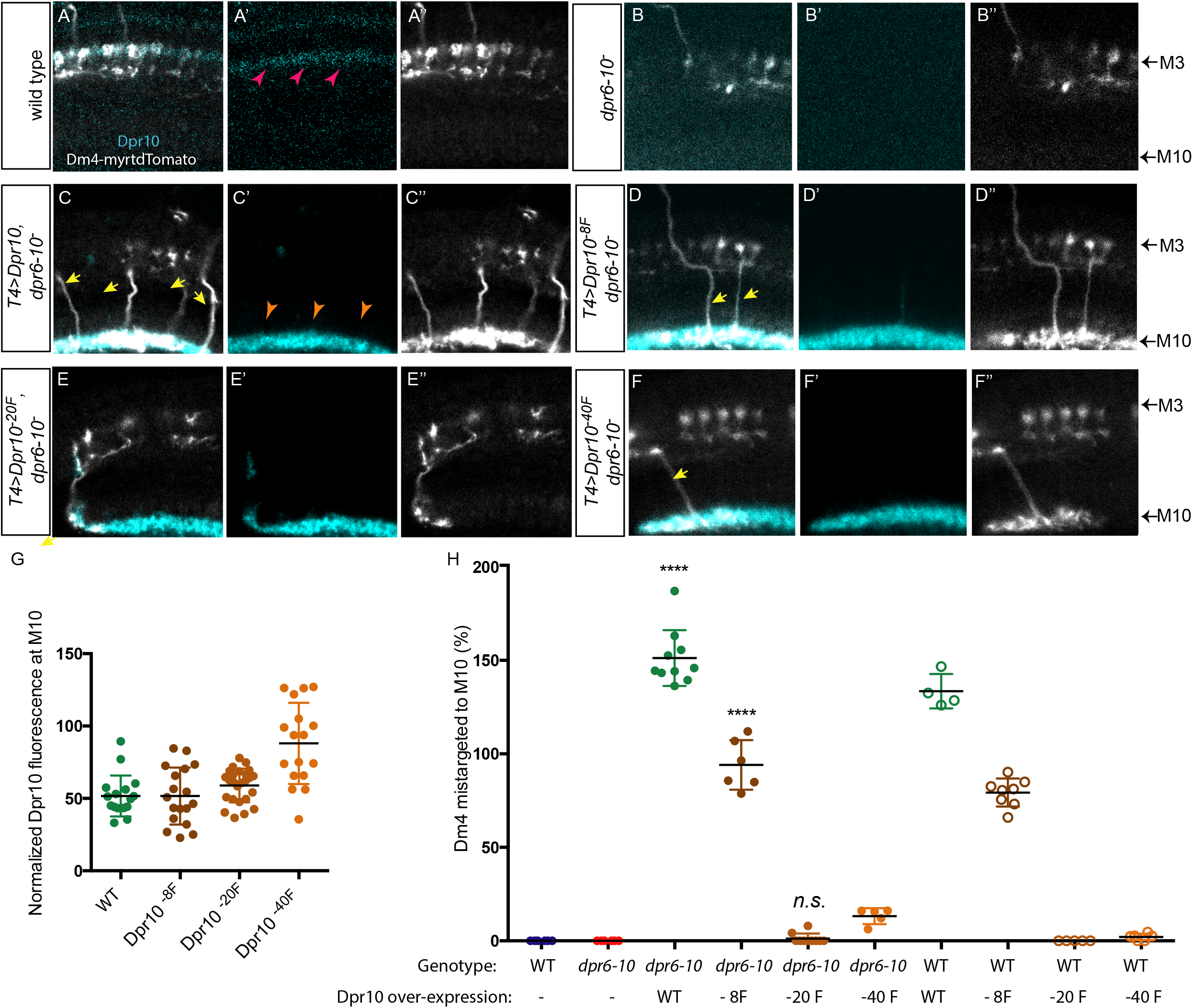
An affinity threshold for induction of mistargeting by ectopic Dpr10. (A-F”) Images showing Dm4 neuron targeting in different genetic backgrounds at 48h APF. A) wild type, B) *dpr6^−^, dpr10^−^*, C) mis-expressing wild type Dpr10 in M10 in *dpr6^−^, dpr10^−^*, D) mis-expressing Dpr10^−8F^ in M10 in *dpr6^−^, dpr10^−^*, E) mis-expressing Dpr10^−20F^ in M10 in *dpr6^−^, dpr10^−^*, F) misexpressing Dpr10^−40F^ in M10 in *dpr6^−^, dpr10^−^*. Images are showing one representative area of a single slice in the whole medulla. G) Normalized fluorescence of anti-Dpr10 antibody staining signals of overexpressed Dpr10 mutant variants in M10 layer. H) Graph summarizing panel A-F, plus data showing over-expression of Dpr10 affinity mutant proteins in the wild-type background. In both genetic backgrounds, Dpr10^−8F^ induces mistargeting but Dpr10^−20F^ does not.

We first analyzed each affinity mutant protein’s ability to induce mistargeting of Dm4 neurons in a *dpr6 dpr10* double null mutant background. In this background, no endogenous Dpr6 or Dpr10 is present in the normal M3 layer (Figures 5A-A’’, pink arrowheads; Figure 5B”). About half of Dm4 neurons undergo cell death due to the lack of Dpr6 and Dpr10, and as a result, there are fewer Dm4 processes in M3. However, the remaining ~20 Dm4 cells are all able to target to the correct layer (Figures 5B-B”). When wild-type Dpr10 was expressed in M10, Dm4 cell death was partially rescued, and most of the Dm4 terminals were attracted to the M10 layer, leaving few terminals in the endogenous M3 layers (Figures 5C-C”). We quantified the axons leaving M3 and targeting to M10 (Figures 5C, D, E, F, yellow arrows), and divided that by the total number of Dm4 neurons in the same sample (counted by cell body number), to calculate the percentage of mistargeted Dm4 neurons. When wild-type Dpr10 was expressed, the percentage of mistargeted Dm4 exceeded 100%, suggesting that some Dm4 neurons send out more than one axon branch to the M10 layer (Figure 5H). Expressing Dpr10^−8F^ in M10 caused about half as much Dm4 mistargeting as wild-type Dpr10, while expressing Dpr10^−20F^ or Dpr10^−40F^ caused almost no mistargeting (Figure 5H).

We also tested the three Dpr10 affinity variants’ abilities to induce Dm4 mistargeting in a wild-type background in which endogenous Dpr6 and Dpr10 are still expressed in M3. In this background, the endogenous Dpr10 in M3 was expected to compete with exogenous Dpr10 in M10, and as a result more Dm4 processes were retained at M3 and fewer mistargeted to M10. As expected, the percentage of mistargeted Dm4 neurons was reduced in all experiments done in this genetic background, but the three variants’ relative ability to induce mistargeting remained the same (Figure 5H). Thus, both LOF data (for DIP-α and Dpr10) and gain-of-function (GOF) data for Dpr10 show that there is a major change in the ability of DIP-α::Dpr10 interactions to direct targeting when the K_D_ of the interaction is reduced from ~11 μM to 28 μM.

### A gradual reduction in cell survival with decreasing DIP-α::Dpr10 affinity is observed for two other types of medulla neurons

We have shown previously that neither Dm4 nor Dm1 exhibits mistargeting in *DIP-α* null or *dpr6, dpr10* double null mutants (Figures 6B, 6F) (Xu et al., 2018). This suggests that DIP-α::Dpr10 interactions are redundant with other cues in directing the processes of Dm4 and Dm1 neurons to the correct layers. On the other hand, survival of Dm4 and Dm1 neurons is affected by null *DIP-α* mutations, so we are able to examine the effects of affinity mutations on cell survival for Dm4 and Dm1, and compare those to the effects observed for Dm12.

**Figure 6:**
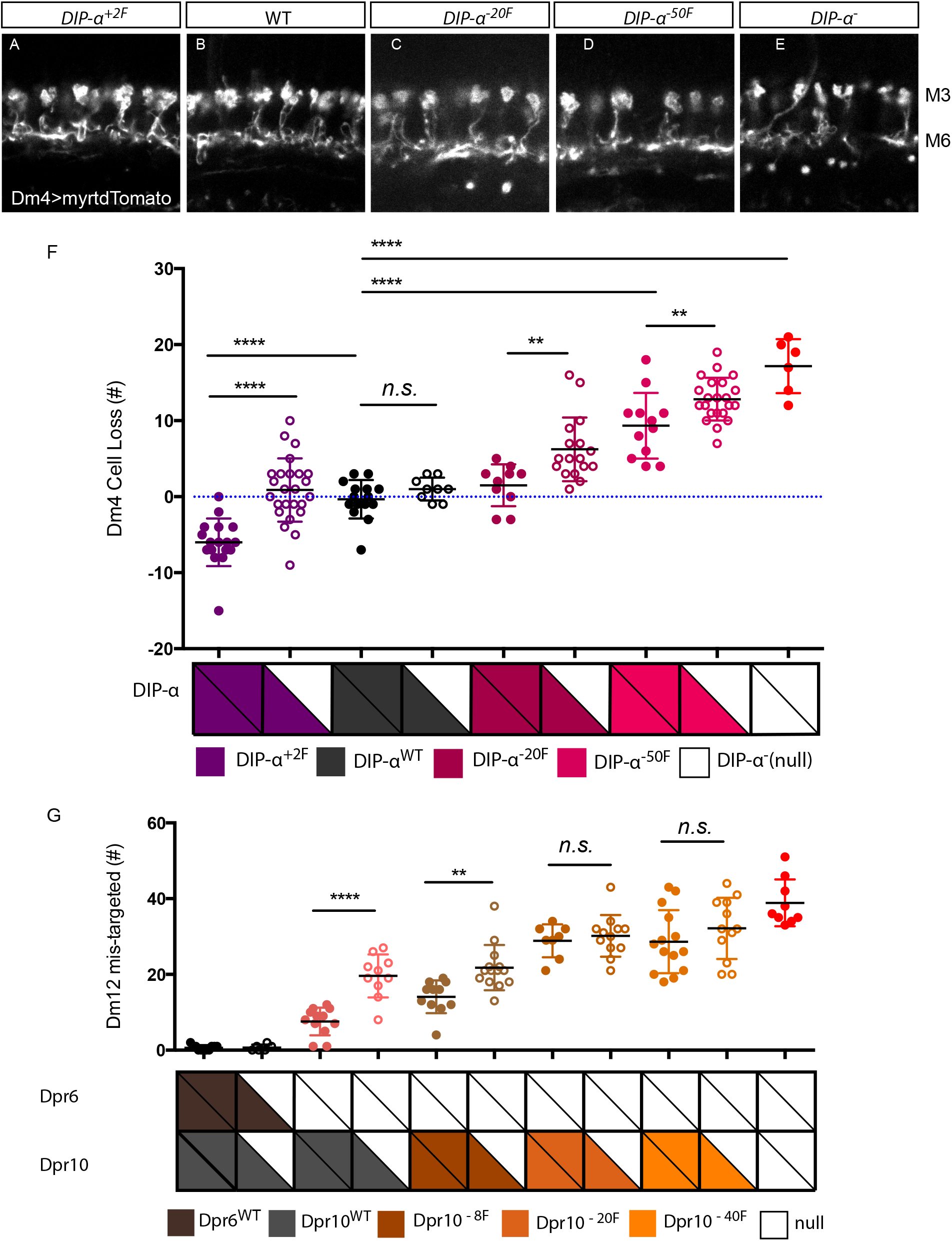
Dm neuron survival and targeting are affected by both individual protein-protein binding affinity and protein copy number. A-E) Dm4 terminals labeled with Dm4-Gal4>UASmyrtdTomato. Genotype, from A to E: *DIP-α^+2F^/DIP-α*, wild type, *DIP-α^−20F^/DIP-α* and *DIP-α^−50F^/DIP-α, DIP-α/DIP-α*,. Images shown are single slices of representative windows within the whole medulla. F) Number of Dm4 cell loss in flies expressing one or two copies of wild type or mutant DIP-α protein. G) Number of Dm12 neurons that mistarget in flies expressing one or two copies of wild type or mutant Dpr6 and Dpr10 protein.

Interestingly, cell survival in Dm4 and Dm1 appears to be more sensitive to affinity reduction than survival of Dm12 neurons. In *DIP-α^−20F^/DIP-α^−^*, cell loss was seen for Dm1 and Dm4, but not for Dm12 (Figures 6C, 6F, Figures S3). The stronger affinity mutant *DIP-α^−50F^/DIP-α-* exhibited as much cell loss as the null allele in Dm1 and Dm4, but had a weaker phenotype than the null in Dm12 (Figures 6D, 6F, Figures S3). These data suggest that different cell types have different affinity thresholds for regulation of cell survival. This could be due to different levels of expression of DIP-α in Dm4 and Dm1 neurons as compared to Dm12, and/or to the use of different cell death pathways. Consistent with the stronger effects of cell loss in *DIP-α* affinity mutants, survival of Dm4 neurons in *dpr10* affinity mutants also displayed a more sensitive response to DIP-α::Dpr10 affinity reduction than survival of Dm12 neurons. Dm4 cell loss is observed in *dpr6 dpr10^−20F^* mutants (Figure S4), which have wild-type numbers of Dm12 neurons (Figure 4).

### Cell surface avidity is a combination of individual protein-protein binding affinity and protein expression levels

The DIP-α^+2F^ mutant protein has a 2-fold increase of binding affinity to Dpr10. In *DIP-α^+2F^/DIP-α^−^* animals, there were no changes in Dm4, Dm12, or Dm1 cell numbers (Figures 3F, 6A, S4). However, when two copies of the same allele were present (*DIP-α^+2F^/DIP-α^+2F^*), we saw a 15% increase in Dm4 numbers (to ~46 cells in adults) (Figure 6F). We have shown previously that Dm4 neurons are produced in excess early during development. Apoptosis happens before 40h APF and eliminates about 20% of the original population, generating the adult number of ~40 Dm4 neurons. If apoptosis is inhibited in Dm4 cells by expressing either the caspase inhibitor baculovirus p35 protein, or the *Drosophila* death-associated inhibitor of apoptosis 1 (Diap1) protein, the number of Dm4 neurons in adults exceeds the number in wild type by 10-15 cells (Xu et al., 2018). Although we knew that the loss of the DIP-α::Dpr10 interaction causes additional Dm4 cell death, it was unknown whether natural apoptosis in Dm4 cells is controlled by DIP-α::Dpr10 interactions. Here, however, we observed that increasing affinity between DIP-α and Dpr10 by only 2 fold can ‘rescue’ ~6 neurons from natural apoptosis.

The difference of Dm4 number between 1 and 2 copies of the mutant gene was also observed in *DIP-α^−20F^/DIP-α^−^ vs. DIP-α^−20F^/DIP-α^−20F^* and *DIP^−50F^/DIP-α^−^ vs. DIP-α^−50F^/DIP-α^−50F^* (Figure 6F). These data suggest that cell surface overall avidity is a combination of protein-protein binding affinity and protein concentration. However, when comparing Dm4 cell number in flies carrying one or two copies of the wild-type *DIP-α* gene *(DIP-α^WT^/DIP-α^−^ vs DIP-α^WT^/DIP-α^WT^*), no changes were observed (Figure 6F).

To examine whether expression levels affect neuronal targeting as well, we analyzed Dm12 mistargeting phenotypes in animals bearing one or two copies of wild-type *dpr10* or the three affinity mutants (Figure 6G). In *dpr6* null animals (bearing two copies of wild type *dpr10*), there are on average ~10 Dm12 neurons that mistarget to M8. Loss of one copy of wild-type *dpr10* in a *dpr6* mutant background produced a significant increase of Dm12 mistargeting, with ~20 Dm12 cells that mistarget to M8 (Figure 6G). Similarly, *dpr6* mutant animals with one copy of *dpr10^−8F^* had stronger phenotypes than those with two copies of *dpr10^−8F^* (22 *vs*. 14 mistargeted axons, respectively) (Figure 6G). However, there were no copy number effects for *dpr10^−20F^* or *dpr10^−40F^* mutations. This is consistent with data in Figure 4, where we showed that there is an affinity threshold for targeting, defined by the transition in phenotypic penetrance observed between ~11 μM (Dpr10^−8F^) and ~28 μM (Dpr10^−20F^). *dpr6 dpr10^−8F^* homozygotes are near that affinity threshold, so removal of ~50% of Dpr10^−8F^ protein by loss of one gene copy had a significant effect. Loss of a copy of *dpr10^−14F^* or *dpr10^−40F^*, however did not affect phenotypic penetrance, presumably because these mutants are below the affinity threshold (Figure 7B). As discussed earlier, *dpr6* mRNAs are expressed at higher levels than *dpr10* (Kurmangaliyev et al., 2020). Removing both copies of *dpr6* already reduced cell surface adhesion. In this background, a small change in Dpr10 affinity significantly affected Dm12 targeting. However, as observed for the wild-type *DIP-α* gene, removing one copy of *dpr6* or *dpr10* in a wild-type background did not result in any Dm12 mistargeting. These results, combined with the finding that Dm4 cell number is not altered by removing one copy of the wild-type *DIP-α* gene (Figure 6F), suggests that wild-type DIP-α::Dpr10 affinity, and likely DIP-α::Dpr6 affinity, is optimal for buffering perturbations in gene expression levels.

**Figure 7.**
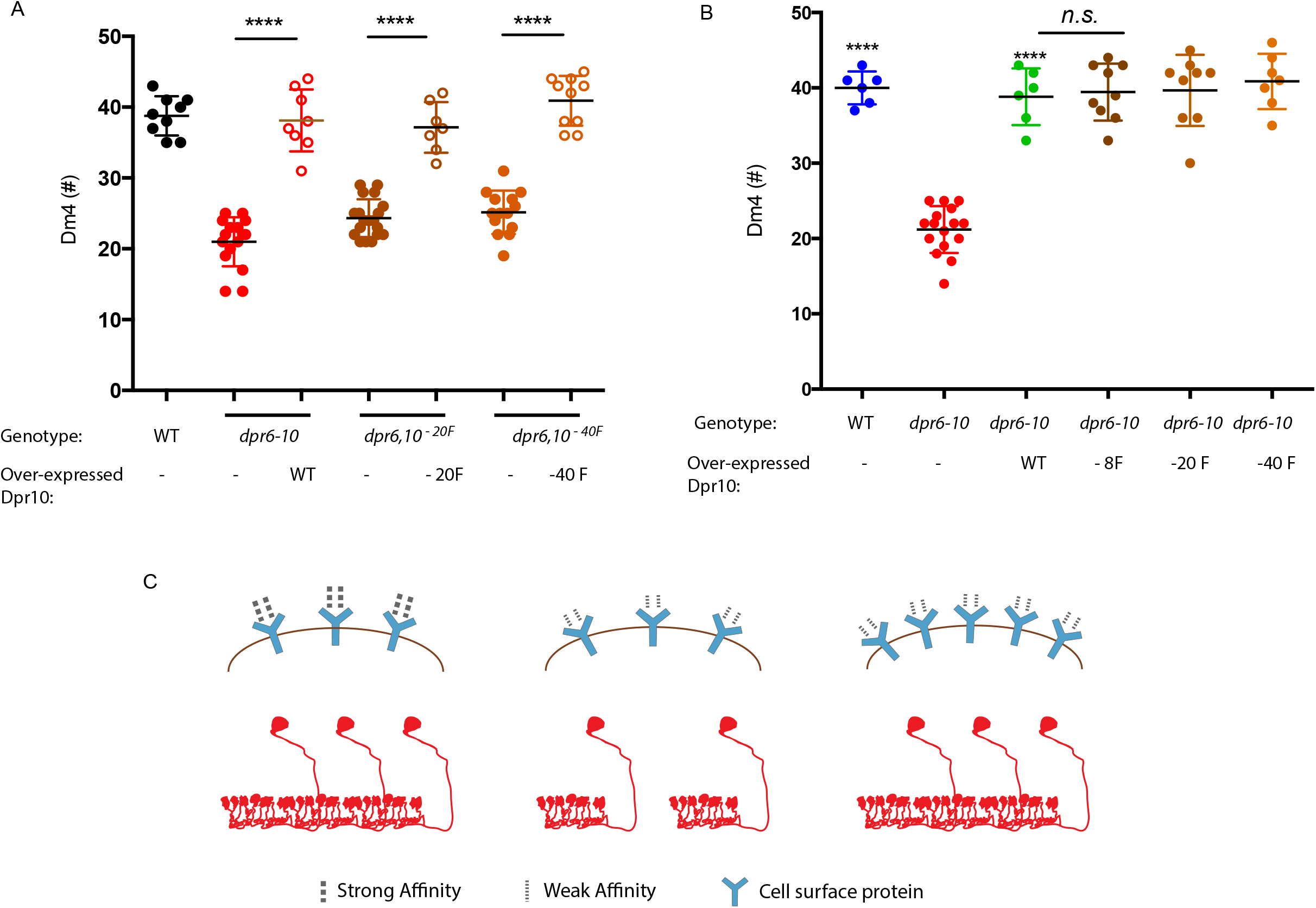
Cell surface avidity is contributed by both individual protein-protein binding affinity and protein expression levels. A) Overexpressing Dpr10^−17F^ or Dpr10^−30F^ in L3 can compensate loss of individual protein-protein binding affinity in *dpr10^−20F^* or *dpr10^−40F^* flies. B) Overexpressing Dpr10^WT^, Dpr10^−8F^, Dpr10^−20F^, or Dpr10^−40F^ in L3 can fully rescue Dm4 cell loss in *dpr6^−^ dpr10^−^*flies. C) Schematic of overexpressing affinity mutant proteins to compensate for reduction in individual protein-protein binding affinity.

### Overexpression of Dpr10 mutants can compensate for reduced protein binding affinity

The results described above indicate that *dpr10* LOF phenotypes can be affected both by altering binding affinity and by changing avidity through alteration of expression levels. To further examine this issue, we asked whether overexpressing Dpr10 affinity mutants could compensate for reduction in individual protein-protein binding affinity. *dpr10^−20F^* and *dpr10^−40F^* mutants both showed significant Dm4 cell loss. We overexpressed these mutant Dpr10 proteins in Dm4’s synaptic partner, the L3 neuron, and analyzed rescue of Dm4 cell loss in the respective mutants. When we overexpressed Dpr10^−20F^ or Dpr10^−40F^ in L3, they fully rescued Dm4 cell loss in *dpr6^−^ dpr10^−20F^* or *dpr6^−^ dpr10^−40F^*, respectively. We previously showed that overexpressing wild-type Dpr10 in L3 can fully rescue Dm4 cell loss in *dpr6^−^ dpr10^−^* double mutants (Figure 7A) (Xu et al., 2018). The mutant proteins Dpr10^−20F^ and Dpr10^−40F^ could fully rescue *dpr6^−^ dpr10^−^* null phenotypes (Figure 7B). These results show that increasing protein amounts can indeed compensate for reductions in individual protein-protein binding affinity (Figure 7C). They also indicate that our affinity mutants would have been classified as fully functional using conventional Gal4 rescue.

Finally, to test whether these results are specific to the Dm4 synaptic partner L3, we overexpressed different UAS-Dpr10 variants in T4 neurons, which project to M10. When wild-type Dpr10 was expressed in T4 cells, it was able to rescue cell loss due to the *dpr6 dpr10* double mutation, but not fully to the wild-type number (34 *vs*. 40). Dpr10^−20F^ or Dpr10^−40F^ were able to rescue Dm4 cell number to similar extents as the wild type Dpr10 protein (Figure S5). The fact that Dm4 neuron survival rescued by T4>Dpr10 is not complete could be due to the fact that T4-Dm4 interaction is transient. Alternatively, there may be other proteins on the L3 surface that contribute to cell survival that are missing from T4 neurons.

## DISCUSSION

In this paper, we systematically explore the impact of CAM affinity and avidity on synaptic connectivity in the *Drosophila* brain, focusing on the Dm4 and Dm12 medulla neurons, which are postsynaptic to the L3 lamina neuron in medulla layer M3. Dm4 and Dm12 express DIP-α, and L3 expresses its binding partners Dpr6 and Dpr10. The loss of interactions between DIP-α and its Dpr partners causes death of Dm4 and Dm12 neurons and mistargeting of Dm12 processes to layer M8. Ectopic expression of Dpr10 in layer M10 redirects Dm4 and Dm12 processes to that layer (Xu et al., 2018).

We generated mutant DIP-α and Dpr10 proteins with altered affinities for each other (Sergeeva et al., 2020). To examine the effects of these affinity alterations *in vivo* without changing expression levels, we introduced the mutations into the endogenous *DIP-α* and *dpr10* genes, so that the mutant proteins would be expressed at endogenous levels (Figure 2). We made fly lines expressing DIP-α mutants that bound to Dpr10 with ~20 and ~50-fold decreases in affinity relative to wild-type, as well as a mutant with a ~2-fold increase in affinity. Dpr10 lines expressed mutants that bound to DIP-α with ~8, ~20, and ~40-fold decreases in affinity (Figure 1).

Both cell survival and targeting are altered by changes in affinity. In Dm12 neurons, mistargeting is more sensitive to affinity reduction than is cell survival. For example, the *dpr10* affinity mutant with a 20-fold reduction in affinity for DIP-α produces a near-null mistargeting phenotype but has no effect on Dm12 cell numbers (Figures 3, 4). Similar affinity requirements are observed for mistargeting of Dm4 processes by expressing ectopic Dpr10: the mutant with an 8-fold decrease in affinity causes mistargeting, but the mutant with a 20-fold decrease does not (Figure 5). We then examined the effects of altering avidity while maintaining affinity by changing the copy number of affinity mutant chromosomes (Figures 6). For both Dm4 cell survival and Dm12 mistargeting, two-fold alterations in copy number have significant phenotypic effects in mutants near the threshold, and having two copies of the *DIP-α* mutant with increased affinity causes more cells to survive than in wild-type. Finally, overexpressing mutants with reduced affinity using GAL4 drivers compensates for the reduction of individual protein-protein binding affinity (Figure 7). This illustrates the importance of introducing mutations into endogenous genes in order to assay affinity effects and other subtle changes in function: increasing avidity through overexpression can allow even very low-affinity mutants to behave like wild-type alleles.

### An affinity threshold for control of Dm12 targeting by DIP-α::Dpr10

In *DIP-α* null or *dpr6 dpr10* double-null mutants, about 1/3 of Dm12 neurons have processes that arborize in layer M8 (Figures 3, 4). Although M8 is distant from M3 in adults, the growth cones of neurons from all medulla layers come into contact with each other during early pupal development, prior to layer separation (Xu et al., 2018). The fact that Dm12 selectively targets M8 suggests that some neurons that arborize in M8 express CAMs that interact with binding partners on the Dm12 surface. Since most Dm12 processes remain in M3 even when DIP-α is absent, however, Dm12 must express other CAMs that allow it to still exhibit a preference for M3 partners. Dm4 neurons also arborize in M3 and are postsynaptic to L3. However, when DIP-α is absent there is no mistargeting of Dm4 processes, suggesting that other CAMs on Dm4 preserve its preference for M3 above any other layers. For both Dm4 and Dm12, these preferences can be overridden by expressing Dpr10 at high levels in neurons that arborize in layer M10. In such cases, most Dm4 and Dm12 processes project to M10 (Figure 5) (Xu et al., 2018).

Reducing Dpr10’s affinity for DIP-α by ~20-fold (from 1.4 μM to 28 μM) produces a near-null phenotype, causing ~30 Dm12 neurons to mistarget to M8. No further increase in mistargeting is seen when affinity is reduced by ~40-fold (Figure 4). This correlates well with the ability of mutant Dpr10 proteins to confer mistargeting of Dm4 processes when they are expressed in M10. Wild-type Dpr10 and Dpr10^−8F^ mutants can redirect Dm4 processes to M10, but Dpr10^−20F^ and Dpr10^−40F^ cannot (Figure 5). These data suggest that there is an affinity threshold between 11 μM and 28 μM.

For DIP-α, we observed that there is substantial Dm12 mistargeting in *DIP-α^−20F^* mutants (K_D_=32 μM), but the phenotype is weaker than that observed in *DIP-α^−50F^* or null mutants (Figure 3). It is also weaker than that for *Dpr10^−20F^*. These data indicate that signaling relevant to synaptic targeting can still occur in response to binding of Dpr10 or Dpr6 to DIP-α proteins with K_D_s of >30 μM. This difference in phenotype between *DIP-α* and *dpr10* mutants with similar affinities may be due to low expression of Dpr10, and high expression of Dpr6, on L3 (Kurmangaliyev et al., 2020; Tan et al., 2015), which could cause a reduction in Dpr10 affinity in a *dpr6* null mutant background to have a stronger effect on the overall avidity of interaction between the Dm12 and L3 cell surfaces than a reduction of the same magnitude in DIP-α affinity.

### Targeting and cell survival have different affinity requirements

Dm4 neurons exhibit cell death during normal development. About 20% of the cells that are originally generated are removed through apoptosis in early pupal development. The Hid pathway controls cell death in Dm4 neurons, while the pathway in Dm12 neurons is unknown. In *DIP-α* or *dpr6 dpr10* double mutants, ~50% of the remaining Dm4 neurons, and about 20% of the Dm12 neurons, undergo apoptosis-mediated cell death during pupal development (Figures 3, 4, 6, S4) (Xu et al., 2018).

Dm12 and Dm4 survival is much less sensitive to alterations in DIP-α::Dpr10 affinity than is Dm12 targeting. No reduction in cell numbers is observed for mutants that reduce affinity of either protein by 20-fold. For both targeting and cell survival, the signaling pathways downstream of DIP-α are likely to be mediated by unknown cell surface proteins, since DIP-α has no cytoplasmic domain (Özkan et al., 2013). Our preliminary analysis suggests that DIP-α is attached to the membrane by a glycosylphosphatidylinositol (GPI) linkage (data not shown). Targeting and cell survival may use different downstream signaling components, accounting for the differential sensitivity of these phenotypes to affinity reduction.

Examination of Dm4 cell number in affinity mutants revealed that increasing binding affinity by 2-fold rescues ~6 cells that would normally die from apoptosis (Figure 6). This indicates that the reductions in cell number that occur during normal development are regulated by DIP-α::Dpr10 interactions. In wild-type animals, signaling through DIP-α may be tuned to facilitate tiling by Dm4s with optimum arbor sizes. *DIP-α* null mutant Dm4 clones in a wild-type background have reduced arbors that cover about 11 medulla columns, vs. 16 columns for wild-type neurons (Xu et al., 2018).

### Relationships between avidity and affinity

We altered avidity while keeping affinity fixed by changing the copy number of affinity mutant chromosomes. We observed that genotypes with one copy of *DIP-α^−20F^* over a null *DIP-α* mutant have significant Dm4 cell loss, while no cell loss is observed for *DIP-α^−20F^/DIP-α^−20F^*. Similarly, removing one copy of *DIP-α^−50F^* enhances the phenotype. Animals with two copies of *DIP-α^+2F^* have decreased cell loss relative to those with two copies of the wild-type allele, but those with one copy of *DIP-α^+2F^* over a null do not (Figure 6).

Alterations in copy number have strong effects on Dm12 mistargeting phenotypes produced by *dpr10* mutations. Loss of *dpr6* alone causes mistargeting of ~8 Dm12 neurons, and this number is doubled when one copy of wild-type *dpr10* is removed in the *dpr6* null background. A similar effect is observed for the *dpr10^−8F^* mutation. However, there is no copy number effect for the *dpr10^−20F^* and *dpr10^−40F^* mutations, which already produce near-null phenotypes when present in two copies (Figure 7). There may be strong avidity effects for *dpr10* because it is present at 10-20 fold lower levels in L3 than *dpr6* in 24-36 hr APF pupae (Kurmangaliyev et al., 2020). Of course, we do not know if these differences in mRNA level correspond to differences in protein abundance. However, it is possible that the *dpr6* null mutation reduces the overall avidity of the L3 cell surface below a threshold required for normal targeting, and loss of one copy of *dpr10* in the *dpr6* background enhances that phenotype, resulting in the mistargeting of ~20 Dm12 neurons.

### Affinity variation in CAM interactions and nervous system assembly

In the immune system, precise binding parameters, including affinities, for interactions between T-cell receptors and their peptide-major histocompatibility complex ligands correlate with different T-cell specificities and activities (for review, see Stone, 2009). In the nervous system, however, the significance of affinity variation among CAMs has not been extensively studied. Given the large number of CAMs that can be expressed by a given neuron, and the ~100 fold variation in affinity among these CAMs, it is likely that the affinities of individual CAM binding pairs have been selected by evolution.

The Dpr-ome has 7 specificity groups, each of which contains one to three DIPs that interact with one to five Dprs (Figure S1) (Carrillo et al., 2015; Cosmanescu et al., 2018; Sergeeva et al., 2020). Segregation into groups is based on protein phylogenetic distance and interaction specificity, which is controlled by “negative constraints” that interfere with binding of non-cognate members (Sergeeva et al., 2020b). The DIP-α group contains DIP-α, Dpr6, and Dpr10, with affinities of DIP-α::Dpr6 and DIP-α:: Dpr10 at 2 μM and 1.36 μM respectively. This is a high affinity interaction group. By contrast, in the DIP-η/θ/Ι and DIP-ε/ζ groups, all K_D_s are >22 μM. Three of the five Dprs in the DIP-η/θ/Ι group (Dprs 1, 3, and 5) have no interactions with K_D_s of less than 70 μM (Figure S1C). Many investigators have studied members of the Dpr-ome in the hopes of finding specific interactions that control synaptic connectivity. In addition to DIP-α interactions with Dpr10 and Dpr6, two other DIP::Dpr interactions have been shown to be essential for brain wiring thus far, DIP-γ::Dpr11, and DIP-δ::Dpr12 (Barish et al., 2018; Bornstein et al., 2019; Carrillo et al., 2015; Courgeon and Desplan, 2019; Menon et al., 2019). These interactions are of relatively high affinity (all K_D_s are <9 μM) (Cosmanescu et al., 2018). The work in this paper shows that high affinity is important for the functions of the DIP-α group. A mutation that changes its K_D_ to 28 μM eliminates the ability of the DIP-α::Dpr10 interaction to drive synaptic targeting (Figures 4, 5).

The fact that interactions within the two low affinity groups all have K_D_s of >20 μM (Figure S1C) raises the question of whether these groups, which include 5 DIPs and 10 Dprs, have different functions than the high affinity groups. The expression patterns of these DIPs and Dprs are similar to those in the high affinity groups, with frequent expression of DIPs in neurons that are postsynaptic to neurons expressing a Dpr that binds to the DIP *in vitro* (Cosmanescu et al., 2018). However, the data obtained thus far has not shown that these DIP::Dpr pairings are required for connectivity. In one study, selectively knocking down DIP-η in the Or47b antennal neurons disrupted the morphology and positioning of the VA1v glomerulus to which these neurons target. However, knocking down DIP-η in all ORNs produced no phenotypes (Barish et al., 2018). In another study, knockdown of DIP-ε in DIP-α-expressing brain neurons produced non-cell-autonomous effects on the morphologies of neurons expressing both Fruitless and DIP-α (Brovero et al., 2020). The above data, while limited, suggest that DIPs in low affinity groups might function differently from those in high affinity groups. It will be interesting to determine whether low affinity is required for such DIPs to function properly, and if increasing affinity for these DIPs will cause defects in wiring. In our future work, we hope to address the mechanisms through which affinity and avidity control CAM signaling for other binding pairs in the OL and other areas of the nervous system.

## EXPERIMENTAL MODEL

Flies were reared at 25°C on standard medium. For developmental analysis and sorting experiments white pre-pupae were collected and incubated for the indicated number of hours. Fly lines used in this study are listed in the Key Resources Table. Genotypes of flies used for each experiment are provided in Supplemental Table 1.

## METHOD DETAILS

### Generation of affinity mutant flies

DIP-α^+2F^, DIP-α^−20F^ and DIP-α^−40F^: The genomic sequence of DIP-α including exon2-exon4 was first replaced with sequence of attP-3XP3-DsRed-attP using a CRISPR-based knock-in strategy (Zhang et al., 2014), generating *DIP-α-PRP* flies. Same above DIP-α genomic region was PCRed out from the wild type fly genome and cloned into pBS-attB vector to make donor plasmid. Mutations (G74A, K81Q, K81Q+G74S) that change DIP-α binding affinity to Dpr10 were introduced into the donor plasmid respectively. Donor plasmids were injected into *DIP-α*-PRP flies generated above. Through ΦC31 recombinase-mediated cassette exchange, mutations were introduced into the fly genome (Injection completed at Bestgene. Inc). Detailed procedure was as in Zhang, *et.al.* 2014.

Dpr10^−8F^, Dpr10^−20F^ Dpr10^−40F^: First, in Vas-Cas9(X);+/+;*dpr6*^null^ flies, the genomic sequence of dpr10 including 2 exons encoding the first Ig domain of dpr10 was replaced with sequence of attP-3XP3-DsRed-attP using a CRISPR-based knock-in strategy (Zhang et al., 2014), generating *dpr6^−^, dpr10-PRP* flies. Same above *dpr10* genomic region was PCRed out from wild type fly genome and cloned into pBS-attB vector to make donor plasmid. Mutations (Q138D, V144K, G99D+Q142E+V144K) that change Dpr10 binding affinity to DIP-α were introduced into the donor plasmid respectively. Donor plasmids were injected into *dpr6^−^, dpr10-PRP* flies generated above. Through ΦC31 recombinase-mediated cassette exchange, mutations were introduced into the fly genome (Injection completed at Bestgene. Inc). Detailed procedure was as in Zhang, *et.al.* 2014.

### Generation of UAS-transgenic flies

cDNA encoding DIP-α-RA and Dpr10-RD were cloned into the pJFRC28 vector (Pfeiffer et al., 2012) using standard cloning methods. V5 sequence was inserted after signal peptide and before Ig1 for DIP-α and Dpr10 as described in Xu, *et. al.* 2018 (Xu et al., 2018). Mutations that change DIP-α and Dpr10 affinity (*DIP-α*: G74A, K81Q, K81Q+G74S; *dpr10*: Q138D, V144K, G99D+Q142E+V144K) were cloned into the above plasmids. Transgenes were inserted into specific landing site at 28E7 by injection of fertilized embryos (Bestgene, Inc.). Plasmid and primer design were carried out using the software Snapgene. Plasmids and detailed sequences are available upon request.

### Immunohistochemistry

Fly brains were dissected in PBS (137mM NaCl, 2.7mM KCl, 10mM Na2HPO4, 1.8mM KH2PO4) and fixed in PBS with 4% paraformaldehide for 30 min at room temperature (RT). After 3 rinses with PBT at RT, samples were incubated in PBT (PBS 0.5% Triton-X10) containing 5% normal goat serum plus 5% normal donkey serum (blocking solution) for at least 1hr at RT. To visualize fine processes of mis-targeted Dm12 neurons, fly brains were fixed in PBT plus 4% paraformaldehyde to increase tissue permeability. Brains were incubated at 4°C in primary and secondary antibodies for at least one day each with multiple PBT rinses at RT in between and afterwards. Brains were mounted in EverBrite mounting medium (Biotium).

The following primary antibodies were used in this study: chicken-anti-GFP (1:1000, Abcam ab13970); rabbit-anti-RFP (1:500, Clontech 600-401-379); mouse-anti-24B10 (Zipursky et al., 1984) (1:20, DSHB); mouse-anti-Brp (nc82) (1:10, DSHB); chicken-anti-V5 (1:500, Abcam 9113); mouse-anti-V5 (1:200, Life Technology, R96025); mouse-anti-DIP-α (4G11) (1:20) (Xu et al., 2018), mouse-anti-Dpr10 (1:5000) (Xu et al., 2018).

Secondary antibodies were used as 1:500 dilution. From Jackson Immuno Research Lab: Alexa Fluor 488 donkey-anti-chicken (703-545-155); Alexa Fluor 488 donkey-anti-mouse (715-545-151); Alexa Fluor 594 donkey-anti-rabbit (711-585-152); From ThermoFisher Scientific: Alexa Fluor 647 goat-anti-mouse (A28181); Alexa Fluor 568 goat-anti-mouse (A-11004). From Life Technologies: Alexa Fluor 647 donkey-anti-mouse (A-21236).

### Microscopy and Image Analysis

Confocal images were acquired on a Zeiss LSM880 confocal microscope. The staining patterns were reproducible between samples. However, some variation on the overall fluorescence signal and noise levels existed between sections and samples. Thus, proper adjustments of laser power, detector gain, and black level settings were made to obtain similar overall fluorescence signals.

### Sedimentation equilibrium by analytical ultracentrifugation

Experiments were performed in a Beckman XL-A/I analytical ultracentrifuge (Beckman-Coulter, Palo Alto CA, USA), utilizing six-cell centerpieces with straight walls, 12 mm path length and sapphire windows. Protein samples were dialyzed to 10 mM Bis-Tris pH 6.6, 150 mM NaCl. The samples were diluted to an absorbance of 0.65, 0.43, and 0.23 at a 10 mm path length and 280 nm wavelength in channels A, B, and C, respectively. Dilution buffer were used as blank. The samples were run at four speeds. Most proteins were run at 16,350, 26,230, 38,440, and 52,980×g. For all runs the lowest speed was held for 20 h and then four scans were taken with a 1 h interval, the second lowest held for 10 h then four scans with a 1 h interval, and the third lowest and highest speed measured as the second lowest speed. Measurements were done at 25 °C, and detection was by UV at 280 nm. Solvent density and protein v-bar were determined using the program SednTerp (Alliance Protein Laboratories). To calculate the K_D_ and apparent molecular weight, data were fit to a global fit model, using HeteroAnalysis software package, obtained from University of Connecticut (http://www.biotech.uconn.edu/auf).

### Surface Plasmon Resonance (SPR) binding experiments

SPR binding assays were performed using a Biacore T100 biosensor equipped with a Series S CM4 sensor chip. To minimize artificial binding resulting from enhanced-avidity effects of oligomers binding to an immobilized ligand surfaces, DIPs are consistently used as ligands and immobilized over independent flow cells using amine-coupling chemistry in HBS pH 7.4 (10mM HEPES, 150mM NaCl) buffer at 25°C using a flow rate of 20 μL/min. Dextran surfaces were activated for 7 minutes using equal volumes of 0.1M NHS(N-Hydroxysuccinimide) and 0.4M EDC(1-Ethyl-3-(3-dimethylaminopropyl)carbodiimide). Each protein of interest was immobilized at ~30μg/mL in 10 mM sodium acetate, pH 5.5 until the desired immobilization level was achieved. The immobilized surface was blocked using a 4-minute injection of 1.0 M ethanolamine, pH 8.5. Typical immobilization levels ranged between 760-980 RU. To minimize nonspecific binding the reference flow cell was blocked by immobilizing BSA in 10 mM sodium acetate, pH 4.25 for 3 minutes using a similar amine-coupling protocol as described above.

Binding analysis was performed at 25°C in a running buffer of 10 mM Tris-HCl, pH 7.2, 150mM NaCl, 1mM EDTA, 1 mg/mL BSA and 0.01% (v/v) Tween-20. Dpr analytes were prepared in running buffer and tested at nine concentrations using a three-fold dilution series ranging from 81-0.012 μM. Similarly, Dpr10 was tested over the DIP-α and DIP-α G74A-immobilized surfaces at eight concentrations using a three-fold dilution series ranging from 27-0.012 μM. Dpr10 Q138D was also tested at a concentration range of 27-0.012 μM over all DIP surfaces, due to a limited protein expression of this mutant. In each experiment, every concentration was tested in duplicate. During a binding cycle, the association phase between each analyte and the immobilized molecule was monitored for either 30 or 40 seconds as indicated by the plotted sensorgrams, followed by 120-second dissociation phase, each at 50 μL/min. At the end of the dissociation phase the signal returned to baseline thus eliminating the need for a regeneration step. The last step was buffer wash injection at 100 μL/min for 60 seconds. The analyte was replaced by buffer every two or three binding cycles to double-reference the binding signals by removing systematic noise and instrument drift. The responses between 25 and 29 seconds, at which point the binding reactions achieve equilibrium as observed by the flat binding responses, were plotted against the concentration of analyte. The data was fit to 1:1 interaction model and the K_D_ was calculated as the analyte concentration that would yield 0.5 R_max_ (Rich and Myszka, 2009). The data was processed using Scrubber 2.0 (BioLogic Software).

## QUANTIFICATION AND STATISTICAL ANALYSIS

Images were analyzed with ImageJ software. Cell number counting were facilitated with Fiji (Schindelin et al., 2012) plugin “ClearVolume” (Royer et al., 2015) and Imaris (Bitplane Inc) software (semi-automatically with hand-correction). Statistial analysis was done using Prism software. All data are shown as mean ± standard deviation (SD). Statistical test: unpaired t-test.

## TRANSGENIC ANIMALS USED IN THIS STUDY

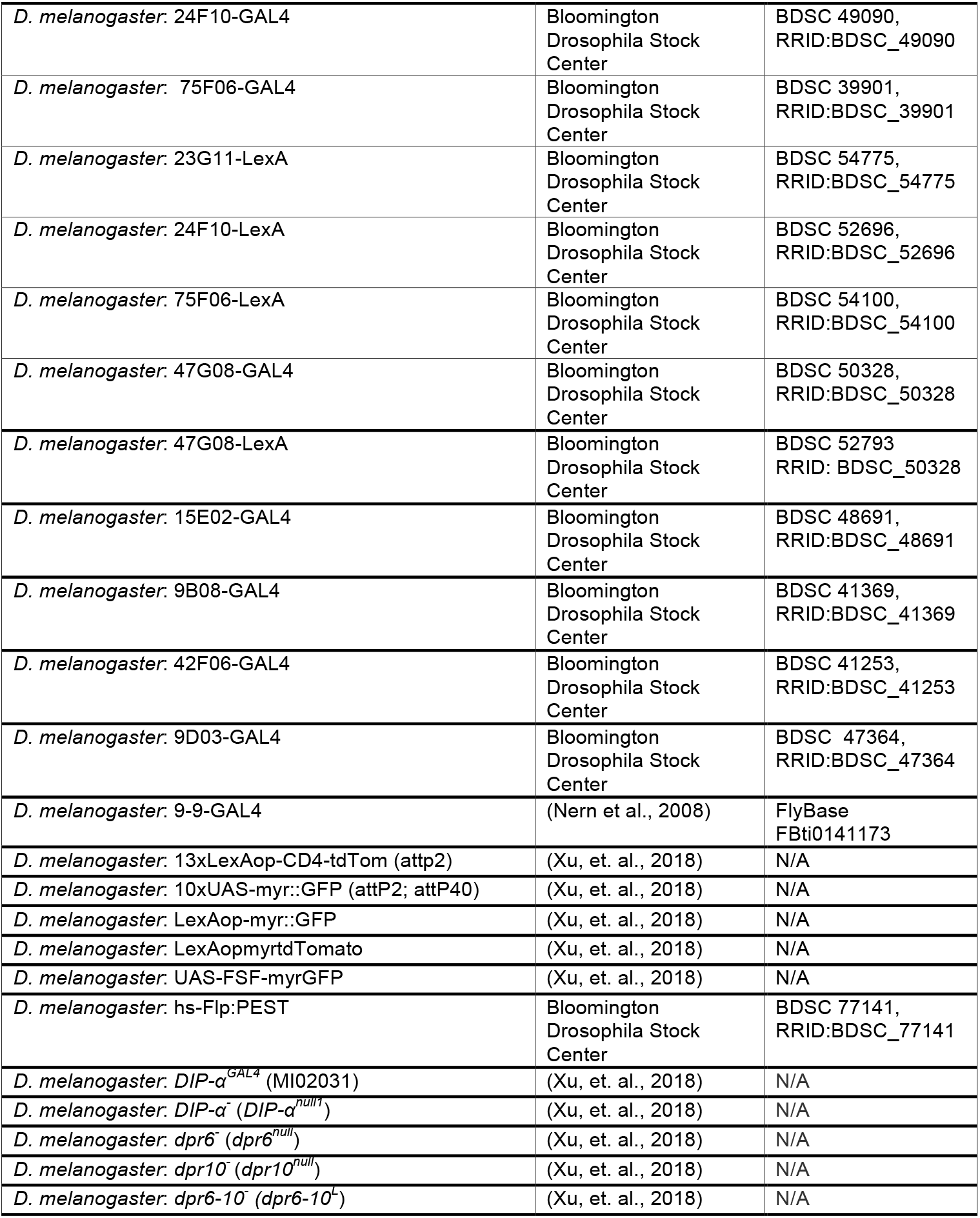
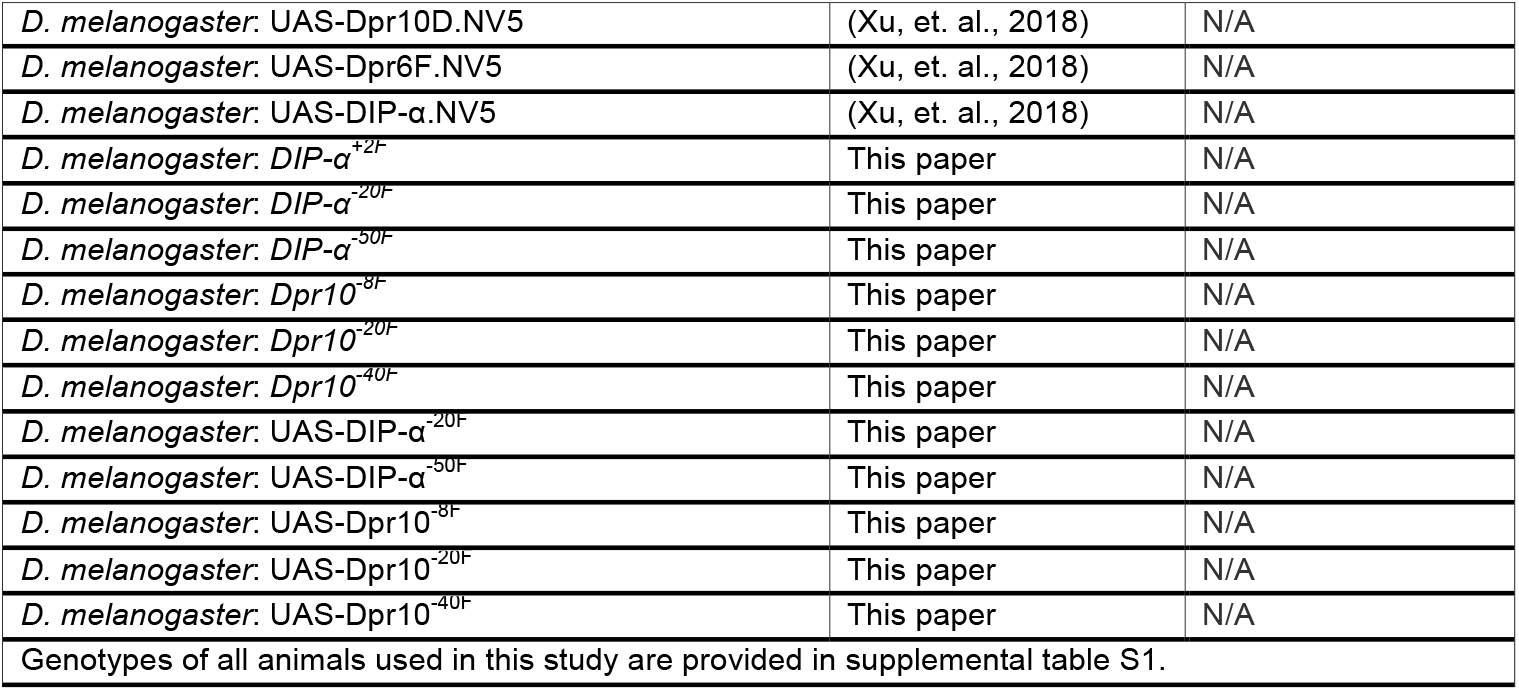

## ACKNOWLEDGEMENTS

We would like to thank Richard Mann, Kaushiki Menon and Namrata Bali for general discussions. Confocal imaging was completed in the Caltech Biological Imaging Facility. This work was supported by Kai Zinn’s NIH grants R37 NS028182 and RO1 NSN096509; and Barry Honig’s NSF grant MCB-1914542.

## Notes

### Competing Interest Statement

The authors have declared no competing interest.

