## Supplemental Information for "Affinity requirements for control of synaptic targeting and neuronal cell survival by heterophilic IgSF cell adhesion molecules"

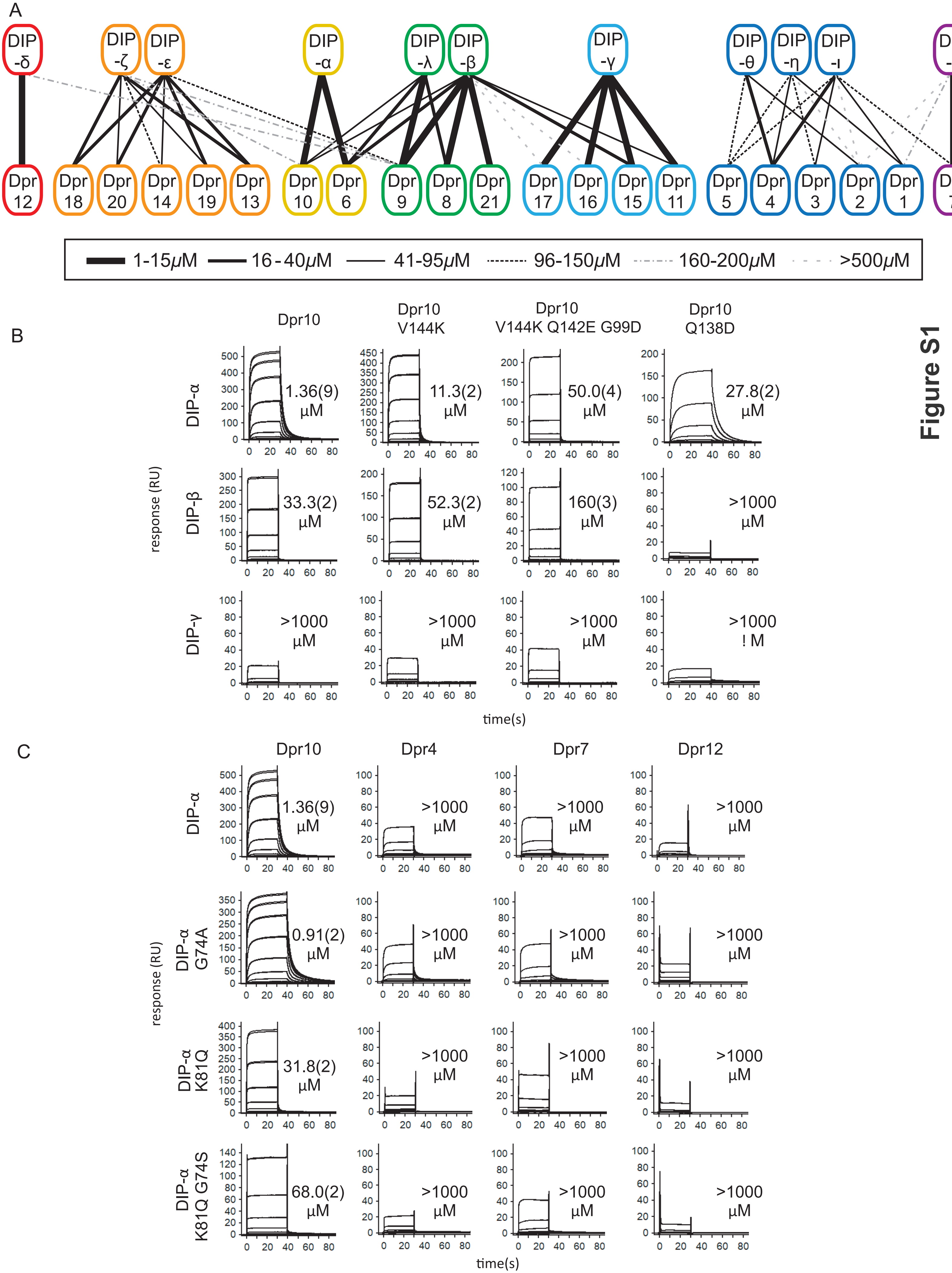
A
DIP -δ
DIP -ζ
DIP -ε
DIP -α
DIP -λ
DIP -β
DIP -γ
DIP -θ
DIP -η
DIP -ι
DIP -κ
Dpr 12
Dpr 18
Dpr 20
Dpr 14
Dpr 19
Dpr 13
Dpr 10
Dpr 6
Dpr 9
Dpr 8
Dpr 21
Dpr 17
Dpr 16
Dpr 15
Dpr 11
Dpr 5
Dpr 4
Dpr 3
Dpr 2
Dpr 1
Dpr 7
1-15μM
16-40μM
41-95μM
96-150μM
160-200μM
>500μM
B
Dpr10
Dpr10 V144K
Dpr10 V144K Q142E G99D
Dpr10 Q138D
DIP-α
DIP-β
DIP-γ
response (RU)
1.36(9) μM
11.3(2) μM
50.0(4) μM
27.8(2) μM
33.3(2) μM
52.3(2) μM
160(3) μM
>1000 μM
>1000 μM
>1000 μM
>1000 μM
>1000 ! M
time(s)
C
Dpr10
Dpr4
Dpr7
Dpr12
DIP-α
DIP-α G74A
DIP-α K81Q
DIP-α K81Q G74S
response (RU)
1.36(9) μM
0.91(2) μM
31.8(2) μM
68.0(2) μM
>1000 μM
time(s)
Figure S1

**Figure S1. SPR binding experiments for Dpr10 and DIP-α mutants**.

(A) Schematic of DIP/Dpr-ome, modified from Sergeeva *et.al* 2020.

(B) SPR responses for Dpr10 and its mutants V144K, V144K Q142E G99D and Q138D, binding over surfaces immobilized with DIP-α (first row), DIP-β (second row), and DIP-γ (third row). Panels, from which accurate $K_D$s could be determined, were plotted on independent scales. The interactions were tested at three-fold dilutions ranging from 0.012-81μM, except for Q138D binding over DIP-α, DIP-β and DIP-γ surfaces, which was tested at a concentration range from 0.012-27 μM.

(C) SPR responses for Dprs 10, 4, 7 and 12 binding over surfaces immobilized with DIP-α (first row), DIP-α G74A (second row), DIP-α K81Q (third row) and DIP-α K81Q G74S (fourth row). Panels for Dpr10, from which accurate $K_D$s could be determined, were plotted on independent scales. These isotherms describing the SPR data are shown in Fig. 1B and C respectively. The interactions were tested at three-fold dilutions ranging from 0.012-81μM, except for Dpr10 binding over DIP-α and DIP-$\alpha^{G74A}$ only, which were tested at a concentration range from 0.012-27 μM.

#### Table S1 (AUC data for homodimer mutants)

| Protein-protein interaction | $K_D$ (homophilic), μM |
|---|---|
| DIP-α/DIP-α WT | 23.9 ± 0.0[a] |
| DIP-α G74A /DIP-α G74A | 50.0 ± 0.6 |
| DIP-α K81Q /DIP-α K81Q | 19.7 ± 1.9 |
| DIP-α K81Q G74S /DIP-α K81Q G74S | 46.3 ± 5.7 |

[a] Published in Cosmanescu et al (2018)

### Figure S2

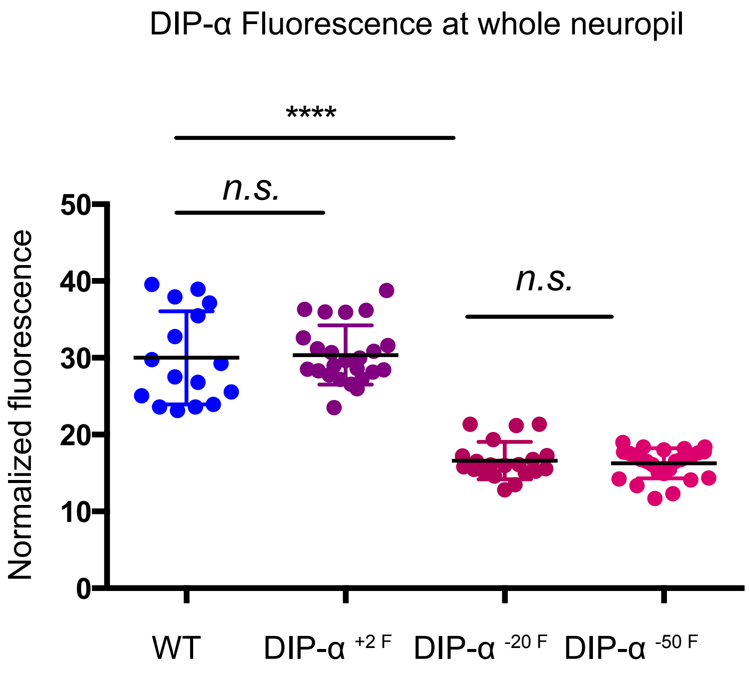
DIP-α Fluorescence at whole neuropil
****
n.s.
n.s.
Normalized fluorescence
50
40
30
20
10
0
WT
DIP-α +2 F
DIP-α -20 F
DIP-α -50 F

**Figure S2**: **anti-DIP-α fluorescence signal intensity of all three medulla layers at 48h APF in wild type and *DIP-α* affinity mutants.**

Fluorescence intensity was normalized against background signal.

### Figure S3

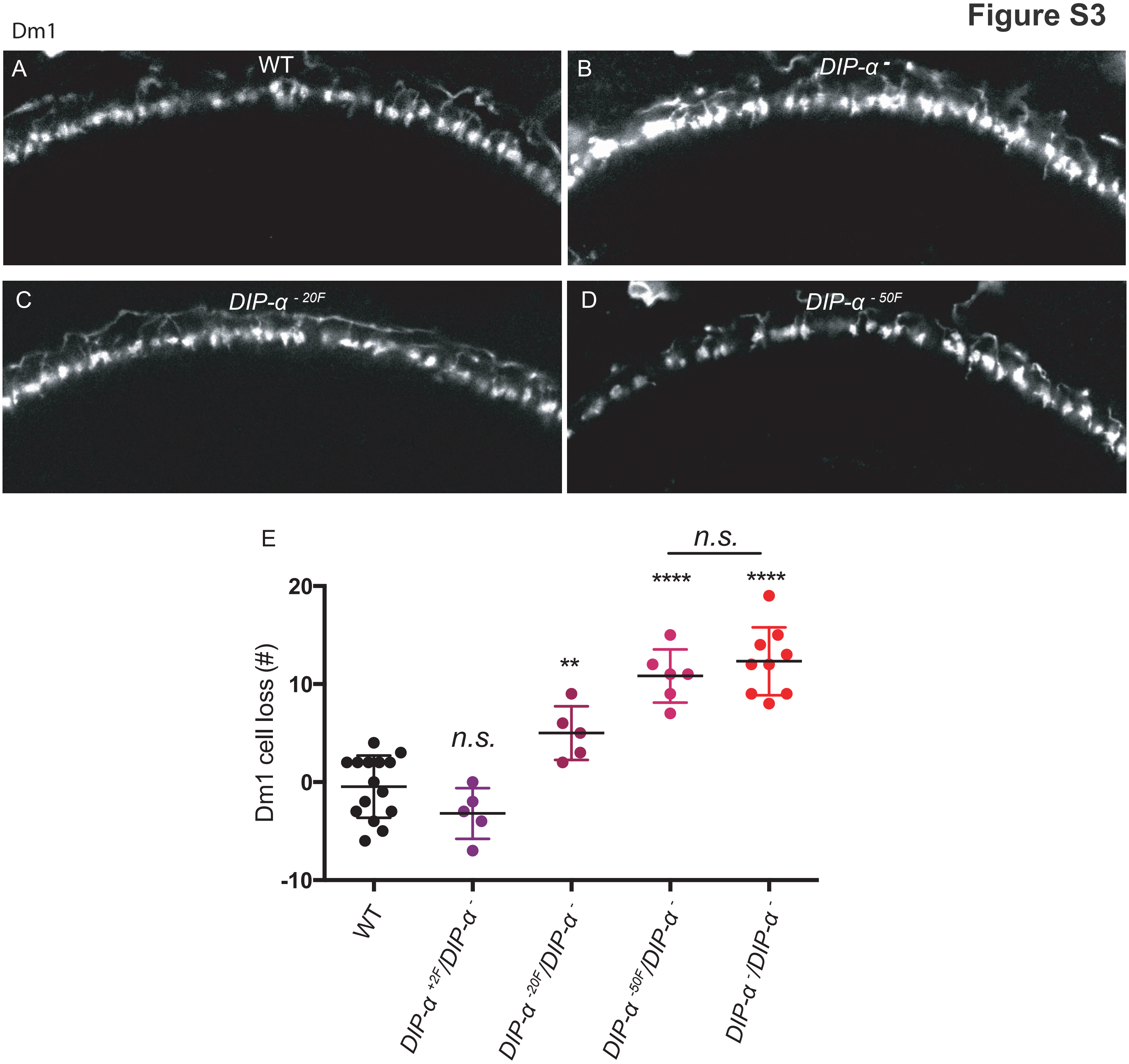

Dm1
A
WT
B
DIP-α -
C
DIP-α - 20F
D
DIP-α - 50F
E
n.s.
****
****
**
n.s.
Dm1 cell loss (#)
20
10
0
-10
WT
DIP-α +2F/DIP-α -
DIP-α -20F/DIP-α -
DIP-α -50F/DIP-α -
DIP-α -/DIP-α -

**Figure S3: Dm1 neurons show gradual cell loss with reduced DIP-α::Dpr10 affinity in *DIP-α* mutants.**

A-D) Dm1 neurons were labeled by Dm1-Gal4;UAS-myrGFP.

E) Number of Dm1 cell lost in different *DIP-α* genotypes.

### Figure S4

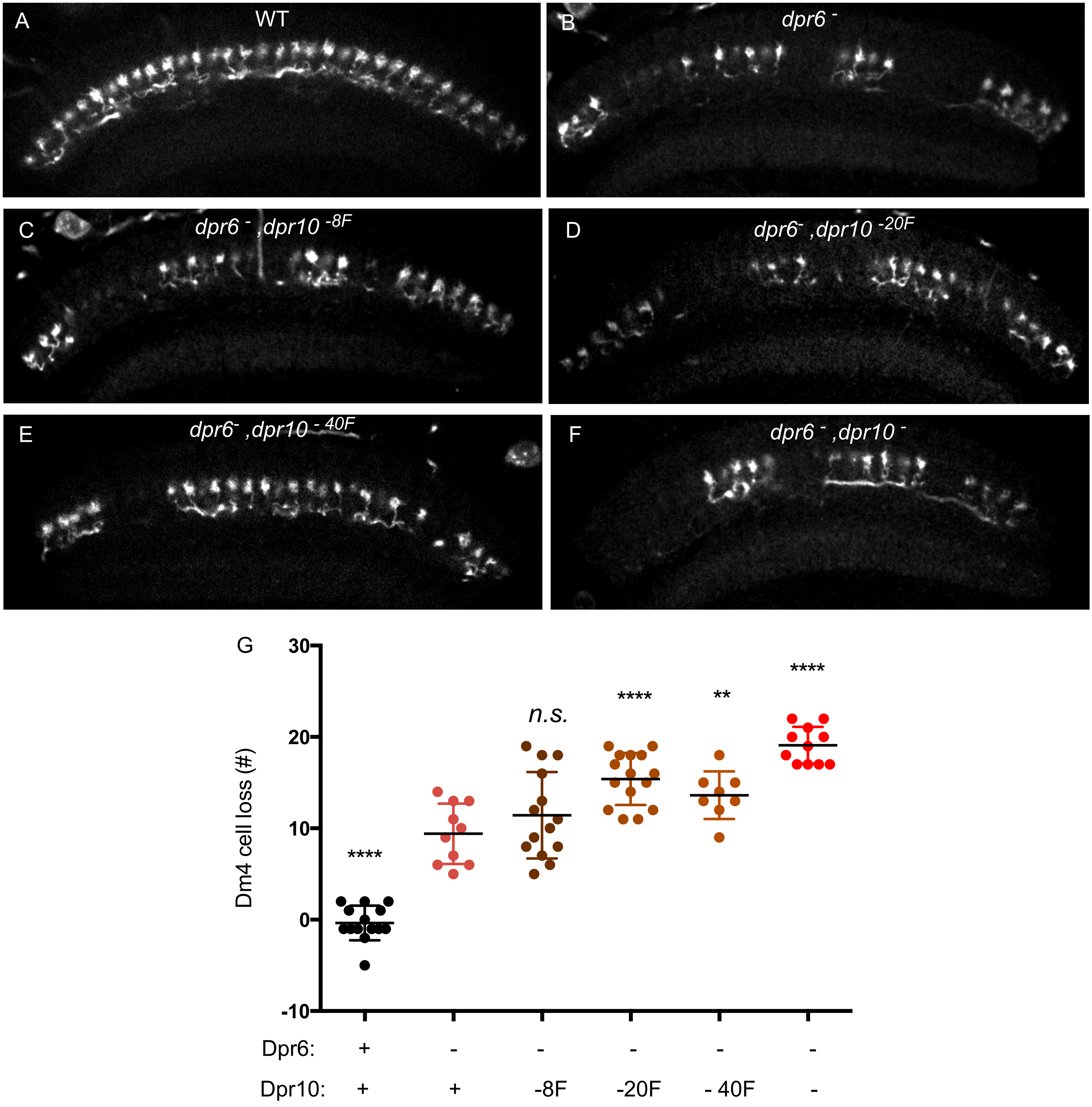

A
WT
B
dpr6 -
C
dpr6 - ,dpr10 -8F
D
dpr6- ,dpr10 -20F
E
dpr6- ,dpr10 - 40F
F
dpr6 - ,dpr10 -
G
Dm4 cell loss (#)
30
20
10
0
-10
****
n.s.
****
**
****
Dpr6: + - - - - -
Dpr10: + + -8F -20F - 40F -

**Figure S4: Dm4 neurons show graded cell loss as DIP-α::Dpr10 affinity is reducing in *Dpr10* mutants.**

A-F) Dm4 neurons were labeled by Dm4-LexA; LexAop-myrtdTomato.

G) Number of Dm4 cell lost in different *dpr10* genotypes.

### Figure S5

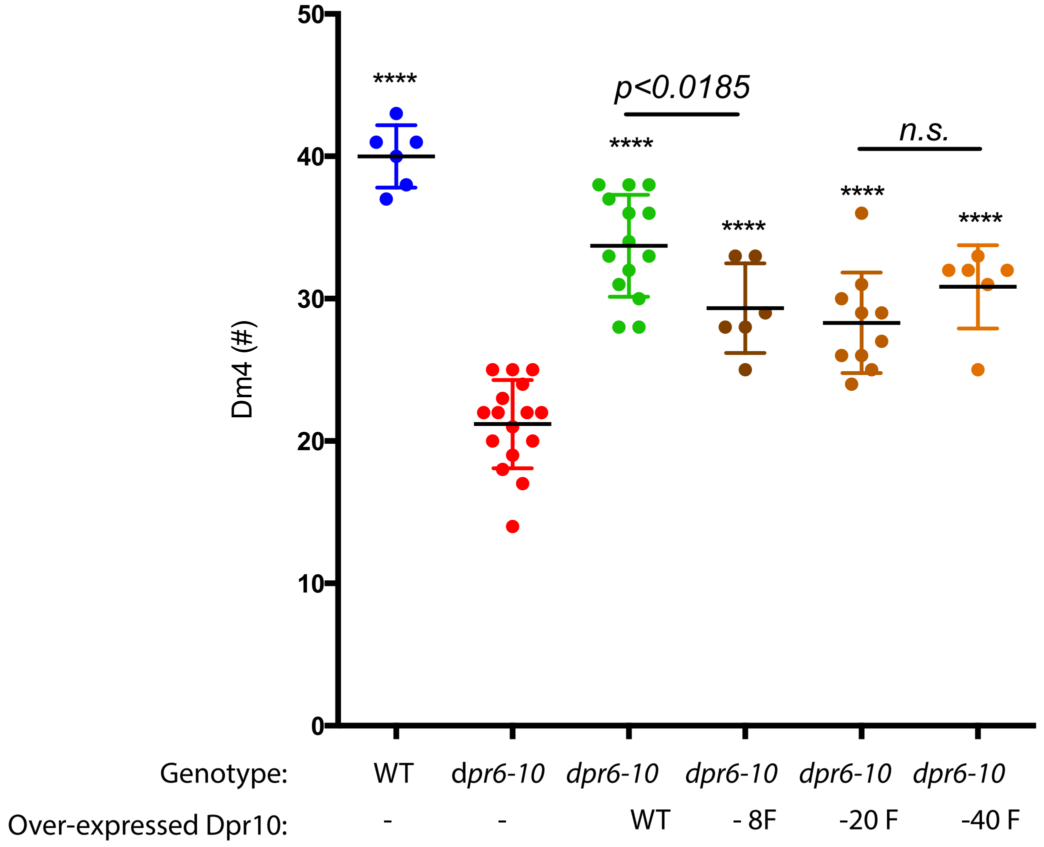
Dm4 (#)
0
10
20
30
40
50
****
****
****
****
****
p<0.0185
n.s.
Genotype:
WT
dpr6-10
dpr6-10
dpr6-10
dpr6-10
dpr6-10
Over-expressed Dpr10:
-
-
WT
- 8F
-20 F
-40 F

**Figure S5**: **Overexpressing Dpr10$^{-8F}$, Dpr10$^{-17F}$ or Dpr10$^{-30F}$ in T4 can partially rescue Dm4 cell loss in *dpr10* flies.**

UAS-Dpr10$^{-8F}$, UAS-Dpr10$^{-20F}$, UAS-Dpr10$^{-40F}$, or UAS-Dpr10$^{WT}$ are expressed by Gal4 driver that drives transgene expression at early pupal development stages before16h APF.
